# Modulation of *rob* expression accelerates development of antibiotic resistance in *Yersinia enterocolitica*

**DOI:** 10.64898/2026.02.23.707304

**Authors:** Xinyu Wang, Taichi Chen, Martijs Jonker, Alphonse de Koster, Wim de Leeuw, Gaurav Dugar, Benno H. ter Kuile

## Abstract

Multidrug-resistant bacteria pose a severe threat to global health. Mutations in transcriptional regulators accelerate the emergence of multidrug resistance and may have a crucial impact on pathogen evolvability under antibiotic exposure. Here, we investigate these dynamics in the bacterial pathogen *Yersinia enterocolitica*. In this organism, we identified a high-frequency *de novo* mutation in the promoter of an AraC/XylS-family transcriptional regulator, Rob. This mutation arose independently during resistance evolution against three of six antibiotic classes. Sequence and structure alignments indicate that Rob is a previously uncharacterized, lineage-specific regulator in *Y. enterocolitica*, featuring a conserved promoter architecture. This promoter mutation resulted in robust *rob* overexpression, leading to the activation of multiple downstream efflux- and membrane-associated pathways. This regulatory mutation emerges early and shapes the acquisition of tetracycline resistance, while also enhancing high-level enrofloxacin resistance in combination with canonical mutations in *gyrA* and *parC*. Despite being favored under antibiotic selection, *rob* overexpression is costly, resulting in counter-selection of the -57 G>A *rob* allele in the absence of antibiotics. Together, these findings identify Rob as a *Yersinia*-specific efflux regulator and demonstrate how regulatory mutations can transiently accelerate antibiotic resistance.

**Graphical abstract:** 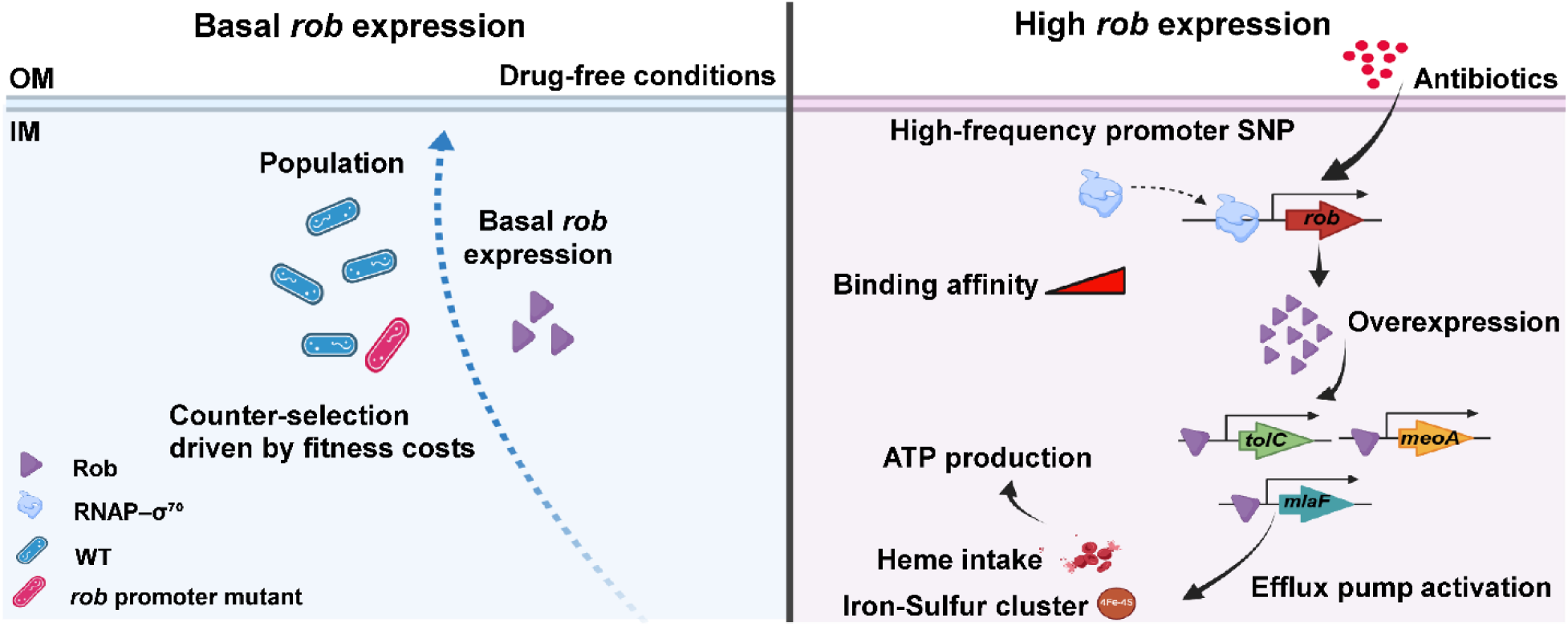

## Introduction

Bacteria rapidly modulate gene expression in response to environmental stress, typically through transient and reversible changes(1–6). In prokaryotes, transcriptional regulators finely tune central metabolic networks to prevailing conditions(7–10), yet this control can be adjusted by single-nucleotide polymorphisms (SNPs) acquired through antimicrobial resistance evolution. Substitutions in the β-subunit of RNA polymerase (RpoB) that disrupt rifampicin binding in *Escherichia coli* convert the drug from bacteriostatic to bactericidal by inducing replication-dependent DNA breaks(11), whereas in *Mycobacterium tuberculosis* the common resistance mutation RpoB (S450L) slows growth through excessive transcriptional pausing that is relieved by compensatory changes in the pausing factor NusG(12). These cases illustrate the importance of mutations in transcriptional regulators in shaping mechanisms of drug resistance.

Members of the AraC/XylS family are prominent regulators of bacterial stress responses, including those in *Salmonella enterica* serovar Typhimurium, *Yersinia pestis*, *Pseudomonas putida*, enteroaggregative *E. coli* and *Shigella* spp(13–16). In *E. coli*, the AraC-like protein Rob binds conserved *marA*/*soxS*/*rob* motifs and activates programs that mediate multidrug resistance(17–19), oxidative-stress defense(20), tolerance to organic solvents(21), and heavy-metal detoxification(22), initiating transcription via a “pre-recruitment” mechanism in which RNA polymerase is engaged before open-complex formation(23). How Rob orchestrates antibiotic responses in *Yersinia enterocolitica* is presently unknown.

Evolutionary studies indicate that regulatory changes often guide the trajectory of resistance. For example, amplification of *sdrM* (a MATE-family multidrug efflux pump) provides an alternative trajectory leading to the development of delafloxacin resistance in methicillin-resistant *Staphylococcus aureus*(24). Overexpression of *norA* (an MFS fluoroquinolone efflux pump) accelerates ciprofloxacin resistance by enhancing the benefit of *gyrA* mutations in the same species(25). Mutations in the two-component systems *pmrB* and *phoQ* constrain the mutational paths to colistin resistance in *Pseudomonas aeruginosa*(26). By analogy, mutations in *rob* may steer resistance evolution in *Y. enterocolitica*, potentially through epistatic interactions with other adaptive changes.

Whole-genome sequencing (WGS) of experimentally evolved *Y. enterocolitica* populations revealed a recurrent mutation in the *rob* promoter that arose independently in the resistance evolution against three of six antibiotic classes. *Y. enterocolitica* encodes two Rob homologs: one that closely resembles the well-characterized Rob protein in *E. coli*, and a second, more divergent, lineage-specific homolog characterized in this study. The Rob protein encoded next to the mutated locus shares only partial sequence and structural similarity with other AraC/XylS family regulators, suggesting it represents a previously uncharacterized variant of this family.

Its strong and repeated association with multidrug resistance highlights the need to define its regulatory function and evolutionary role. Here, we investigate how a regulatory mutation upstream of *rob* influences its expression, examine the genome-wide consequences of altered Rob activity, and evaluate how Rob modulation shapes the evolutionary dynamics of antimicrobial resistance in *Y. enterocolitica*.

## Materials and Methods

### Bacterial Strains and Reagents

*Yersinia enterocolitica* DSM 4780 was purchased from DSMZ (German Collection of Microorganisms and Cell Cultures). All experiments were conducted in Tryptic Soy Broth (TSB). Bacteria from a -80 °C glycerol stock were streaked onto Tryptic Soy Agar (TSA) plates. The *Y. enterocolitica* strains were cultured at 30 °C under aerobic conditions to ensure sufficient growth. Antibiotic solutions, including enrofloxacin (ENR), tetracycline (TET), amoxicillin (AMO), erythromycin (ERY), chloramphenicol (CHL), cefepime (CEP), cefotaxime (CTX), fosfomycin (FOS), rifampicin (RIF), and kanamycin (KAN) were prepared from powder stocks (Sigma, Germany) and filtered through a 0.22 µm pore-size filter (Merck, USA). Antibiotic stock solutions were stored at -20 °C until use. Amoxicillin solutions were freshly prepared and stored at 4 °C for a maximum of four days.

### Rob Protein Evolutionary Analysis and Structure Mapping to MarA-DNA complex

Sequences used for Rob evolutionary analysis were obtained from a strain-defined collection of AraC-family homologous proteins, as provided in the FASTA file: *Escherichia coli* K-12 (MarA, SoxS, Rob), *Yersinia enterocolitica 8081* (Rob_1, Rob_2), *Yersinia pestis C092* (Rob), *Yersinia pseudotuberculosis IP32953* (Rob), *Salmonella enterica* serovar Typhimurium LT2 (Rob, RamA), *Klebsiella pneumoniae* NCTC13443 (RamA) and strain 295R (Rob), *Enterobacter cloacae* E1252 (Rob), *Citrobacter freundii* ICC168 (Rob), and *Shigella flexneri* 301Ser2a (Rob). All protein sequences used in this study are provided in the Supplementary Data. Protein multiple-sequence alignment was performed with Clustal Omega (default settings), and phylogenetic inference was conducted on the resulting alignment using NGPhylogeny.fr. For structural mapping, the AlphaFold2-predicted model of *Y. enterocolitica* Rob_1 was structurally aligned with the *E. coli* MarA–DNA complex (**PDB: 1BL0**), which served as a reference for mapping the helix–turn–helix DNA-binding region using RCSB PDB structural alignment.

### Mutant Construction

The *Yersinia enterocolitica* DSM 4780 strain carrying the *rob* -57 G>A promoter mutation was generated via homologous recombination. Compared with the parental DSM 4780 strain, this mutant exhibits increased *rob* expression and is therefore designated the *rob* overexpression strain (DSM 4780 *rob* -57 G>A). A DNA fragment encompassing approximately 500 bp upstream and downstream of the single-nucleotide polymorphism was amplified by PCR from a mutant isolated during the evolutionary assay using specific oligonucleotides (**Table S1**). The amplified fragment was cloned into the suicide vector pDS132 using the ClonExpress Ultra One Step Cloning Kit (Vazyme) and transformed into the conjugative *Escherichia coli* S17-1 strain. The construct was subsequently introduced into wild-type *Y. enterocolitica* by conjugation, and mutant selection was performed on *Yersinia*-selective agar plates containing 17 µg/mL chloramphenicol. To generate the Rob-Flag-tagged strain, a C-terminal 3xFLAG tag was inserted at the native *rob* locus using the same recombination strategy.

### Antimicrobial Susceptibility Testing

Minimum inhibitory concentration (MIC) assays were performed using twofold serial dilutions of antibiotics in TSB(27). Each measurement was conducted from an overnight culture, in a 96-well plate. Absorbance was measured at 595 nm using a Thermo Scientific Multiskan FC reader with SkanIt software. Each well contained 150 µL of TSB inoculated with overnight culture to a starting OD_595_ of 0.05. Measurements were taken after 24 hours of incubation at 30 °C. A cut-off OD_595_ value of 0.2 was used to define resistance, and growth curves were monitored to validate the MIC determinations.

### Antimicrobial Resistance Evolution

The evolution experiments for resistance development were initiated from single colonies of the wild-type and *rob* mutant strains(27). Evolution was conducted by exposing four replicates of the strains to step-wise increasing sublethal concentrations of enrofloxacin and tetracycline. Three replicates each of the wild-type and *rob* mutant strains grown without antibiotics, served as biological controls. Bacteria were isolated from TSA plates and cultured in 5 mL of TSB in tubes, incubated with shaking at 220 rpm for 24 hours per cycle. Strains were initially exposed to 1/8 MIC of tetracycline or 1/16 MIC of enrofloxacin, with an inoculum size adjusted to achieve a final OD_595_ of 0.1 (4.52 × 10⁸ CFU/mL results in OD_595_ = 1). Antibiotic concentrations were doubled when incubations grew more than 75% of the growth observed in the biological control, based on OD measurements. The experiments continued until strains developed stable, high-level resistance. Stocks of evolved populations were stored at −80 °C in 30% glycerol.

### Gene Expression Analysis by Quantitative PCR (qPCR)

RNA from wild-type and *rob* mutant strains was isolated following the Quick-Start Protocol of the Qiagen RNAprotect® Bacteria Reagent. RNA concentrations were measured using a NanoDrop 2000/2000c spectrophotometer (Thermo Scientific™). A concentration of 35 ng/µL RNA was used for cDNA synthesis. The cDNA reaction was run in a PCR machine with the following program: 25 °C for 5 minutes, 46 °C for 20 minutes, and 95 °C for 1 minute. Quantitative PCR (qPCR) was then performed to measure the expression levels of the housekeeping gene *16S rRNA* and the *rob* gene (**Table S1**). Each qPCR reaction contained 0.5 µM forward primer, 0.5 µM reverse primer, and SYBR Green (2×). For each well, 9 µL of qPCR mix and 1 µL of cDNA were combined. The qPCR was run on an Applied Biosystems 7300 Real-Time PCR System.

### Electrophoretic Mobility Shift Assay (EMSA)

*rob* promoter sequences (150 bp, 10 nM) from wild-type and *rob* mutant strains (**Table S1**) were co-incubated with *E. coli* RNA polymerase holoenzyme containing either RpoD or RpoS (New England Biolabs) at final concentrations of 60 nM, 40 nM, 20 nM, 10 nM, and 0 nM for 30 minutes at 30 °C. Reactions were performed in binding buffer consisting of 200 mM Tris-HCl (pH 7.5), 192 mM KCl, 25 mM MgCl₂, 25% (v/v) glycerol, and 0.5 mM EDTA. Reaction mixtures were separated on a 6% native PAGE gel, and DNA bands were visualized using SYBR Gold nucleic acid stain (Thermo Fisher Scientific).

### Measurement of Maximum Exponential Growth Rate

The growth rates of wild-type and *rob* mutant strains exposed to 1/16 MIC of enrofloxacin, 1/8 MIC of tetracycline, and drug-free conditions were determined using a plate reader (Thermo Scientific, Multiskan FC). Overnight cultures were diluted to an OD_595_ of 0.05 in 150 µL TSB, and OD measurements were initiated immediately. The plates were incubated in the plate reader for 24 hours, with OD_595_ measurements recorded at 10-minute intervals. Growth rates were analyzed in RStudio using a script downloaded from GitHub (https://github.com/Pimutje/Growthrates-in-R/releases/tag/Growthrates). The significance of growth rate differences between resistant strains and controls was assessed using a t-test, with a p-value ≤ 0.05 considered statistically significant.

### Efflux Efficiency Measurement

Efflux pump activity was determined following a previously described protocol(28, 29). Single colonies of wild-type and *rob* mutant strains were grown in 5 mL of TSB at 30 °C with shaking at 200 rpm for 23 hours. Cultures were incubated with carbonyl cyanide m-chlorophenyl hydrazone (CCCP) at a final concentration of 5 µM for 15 minutes at room temperature. Subsequently, either no drug (drug-free control), 1/16 MIC of enrofloxacin, or 1/8 MIC of tetracycline was added, followed by an additional 15-minute incubation at room temperature. Nile Red was then added to a final concentration of 5 µM, and cultures were incubated for 3 hours at 30 °C with shaking at 200 rpm. After incubation, 150 µL of each cell suspension was transferred to a 96-well plate. Immediately before fluorescence measurement, 50 µL of filter-sterilized glucose was added to each well to a final concentration of 25 mM. Fluorescence was measured from the top of the wells using excitation and emission wavelengths of 550 nm and 650 nm, respectively. For quantitative comparisons, fluorescence values were recorded at the final time point of the assay (cycle 15, approximately 11 min after glucose addition), representing endpoint fluorescence measurements.

### Genomic DNA Isolation from Colonies and Populations

In total, genomic DNA was isolated from 80 evolved populations and 6 medium-adapted populations using the PureLink Genomic DNA Kit (USA). Wild-type and *rob* mutant strains exposed to tetracycline at days 4, 8, 12, and 18, with four biological replicates at each time point, were selected for sequencing. Similarly, wild-type and *rob* mutant strains exposed to enrofloxacin at days 3, 6, 9, 12, 15, and 20 were also chosen. Additionally, three samples each of the wild-type and *rob* mutant strains grown without antibiotic exposure were sequenced to control for mutations not associated with antimicrobial resistance.

During drug-free adaptation, individual colonies were isolated from evolving populations between days 3 and 5 of passaging. Fourteen single colonies were selected for analysis, including isolates carrying either the -57 G>A promoter mutation or the wild-type nucleotide at this position. Genomic DNA was extracted from each isolate and subjected to WGS to characterize their genomic backgrounds.

### Whole-Genome Sequencing (WGS)

Genomic DNA libraries were prepared using the NEBNext Ultra II FS DNA Library Prep Kit for Illumina (New England BioLabs), targeting an insert size distribution of 275–475 bp based on the corresponding size selection option provided in the protocol. The libraries were then clustered and sequenced (2 × 150 bp paired-end) on a NextSeq 550 Sequencing System (Illumina) using a NextSeq 500/550 Mid Output v2.5 kit (300 cycles; Illumina). Sequencing coverage depth was set to target an average of 120× to 170× per sample. Raw reads were trimmed and aligned to the reference genome GCA_901472495.1 (*Y. enterocolitica*) using Bowtie2. Variant calling and downstream analyses were performed as described below(30).

### Colony Isolation and Sanger Sequencing

Forty single colonies were isolated from enrofloxacin-evolved populations of the wild-type strain collected between days 9 and 15. The resistance gene alleles of *gyrA*, *gyrB*, *parC*, and *rob* were amplified using PCR with gene-specific primers (**Table S1**) and Rapid Taq Master Mix (Vazyme). PCR products were purified with the Invitek Molecular MSB® Spin PCRapace Kit. Sanger sequencing was then performed to identify the mutations present in each colony.

### Haplotype analysis of evolved populations

Haplotypes in evolved populations were determined by combining analyses of both population sequencing data and isolated colonies. For mutations within the same gene, IGV was used to confirm if multiple mutations co-occurred within single colonies by analyzing single reads covering both mutation sites. For mutations occurring in different genes, haplotypes were identified by isolating 40 single colonies from evolved populations and verifying their mutational profiles using Sanger sequencing. Additionally, changes in allele frequency dynamics over time provided insights into genotype composition and evolution trajectories.

### RNA Isolation and Transcriptome Sequencing

Total RNA was isolated from mid-logarithmic growth phase of cells exposed to the following conditions: drug-free, 1/16 MIC of enrofloxacin, 1/8 MIC of tetracycline. Under these conditions, the growth of both wild-type and *rob* mutant strains were minimally affected by the sub-lethal antibiotic concentrations. RNA isolations were performed in three biological replicates using the Qiagen Quick Start protocols for the RNAprotect® Bacteria Reagent, RNeasy® Protect Bacteria Kit, and RNeasy® Mini Kit. RNA concentrations were measured using a NanoDrop 2000/2000c Spectrophotometer (Thermo Scientific™). Samples were normalized to 200 ng/µL and submitted to the Dutch Genomics Service & Support Provider (University of Amsterdam), where RNA-seq libraries were prepared using the NEBNext rRNA Depletion Kit (Bacteria) in combination with the NEBNext Ultra II Directional RNA Library Prep Kit for Illumina(31). Sequencing libraries were prepared and sequenced on a NextSeq 550 Sequencing System (Illumina) with 75 bp read lengths. Raw RNA-seq reads were quality-trimmed and aligned to the *Yersinia enterocolitica* reference genome (GCA_901472495.1) using Bowtie2 on the Galaxy platform. Differential gene expression analysis was performed with DESeq2, which applies Benjamini–Hochberg correction for multiple testing. Normalized gene expression changes (log₂ fold changes) were calculated by comparing resistant strains to antibiotic-free controls, and genes with an absolute fold change ≥ 2 and an adjusted *P* value (FDR) < 0.05 were considered differentially expressed. Functional enrichment analysis was conducted based on COG annotations.

### Chromatin Immunoprecipitation Sequencing (ChIP-seq) and analysis

Rob-Flag-tagged and wild-type strains, each with three biological replicates, were exposed to 1/8 MIC of tetracycline and grown in 20 mL flasks for 4 hours. Followed by transcription initiation blocking with rifampicin (final 150 mg/L; stock 50 mg/mL in methanol, 300 µL per sample) for 20 min at 28 °C, 220 rpm; cells were crosslinked by adding 37% formaldehyde to 1% final for 20 min at 28 °C, 220 rpm, and the reaction was quenched by adding 2.5 M glycine to 125 mM final, incubating 20 min at 4 °C with gentle shaking; cells were pelleted (6000×g, 5 min, 25 °C), washed three times with 1× PBS, and pellets were processed immediately or stored at −80 °C; pellets were resuspended in 200 µL lysozyme (20 mg/mL) and incubated 30 min at 37 °C, chromatin was fragmented by sonication (Covaris, M220 Focused-ultra-sonicator) for 3 minutes to generate DNA fragments of approximately 150–400 bp, debris were removed (14,000 rpm, 10 min); lysates were pre-cleared with 50 µL Protein A/G beads (MedChemExpress) for 2 h at 4 °C on a rotating wheel, beads were pelleted (13,800×g, 1 min) and the supernatant was incubated with 10 µL mouse anti-FLAG antibody overnight at 4 °C with rotation, then immune complexes were captured with 50 µL Protein A/G beads for 6 h at 4 °C (or overnight) and beads were collected (10,000 rpm, 5 min), washed four times with lysis buffer (NaCl, HEPES, sodium deoxycholate, Triton X-100, DTT, protease inhibitor cocktail 1:100, and Tris-HCl) at 4 °C (1 mL each; 14,000 rpm, 5 min) and twice with TE buffer; complexes were eluted with 100 µL elution buffer 1 at 65 °C for 10 min and then 150 µL elution buffer 2, pooling eluates (250 µL total), and crosslinks were reversed at 65 °C for 6 h or overnight; 10% eluate (25 µL) was reserved for optional Western blot, the remainder was treated with RNase A (20 µL; 100 mg/L) for 30 min at 37 °C and digested with proteinase K (250 µL of a proteinase K/glycogen mix prepared in TE) for 2 h at 58 °C, followed by DNA cleanup with 4 M LiCl (55 µL), phenol:chloroform:isoamyl alcohol extraction (25:24:1) and ethanol precipitation (−80 °C, 30 min; 14,000 rpm, 5 min, 4 °C), washing with 70% ethanol, air-drying, and resuspension in 25 µL nuclease-free water; DNA was quantified (Qubit) and stored at −20 °C.

Immunoprecipitated DNA was subjected to ChIP-seq library preparation using Novogene’s standard workflow, including end repair, A-tailing, Illumina adapter ligation, size selection, and PCR amplification, and sequenced by Novogene (Munich, Germany) on an Illumina NovaSeq X Plus platform with paired-end 150 bp reads. Raw sequencing reads were trimmed and aligned to the *Yersinia enterocolitica* reference genome (GCA_901472495.1) using Bowtie2. DNA-binding enrichment was visualized using IGV. ChIP-seq peaks were identified using MACS2. Aligned and sorted BAM files from ChIP samples were jointly used as treatment, with corresponding control samples provided as input. Peak calling was performed in BAM mode with a genome size of 4.6 × 10⁶ bp. Motif discovery was performed using MEME-ChIP on the eight peaks identified using MACS2.

### Amplicon Sequencing

In total, 15 genomic DNA samples were isolated at days 3, 5, 7, 10, and 15 from adapted populations of three biological replicates of the *rob* mutant strain without antibiotic exposure. A total of 1.5 µg of genomic DNA was used as the template to amplify the *rob* allele locus by PCR (**Table S1**). PCR products were purified using the Invitek Molecular MSB® Spin PCRapace Kit. Amplicon samples were quantified using the TapeStation D5000 ScreenTape system (Agilent). A total of 200 fmol of each amplicon was used as input material for the Native Barcoding Kit 24 V14 (SQK-NBD114-24, Oxford Nanopore Technologies), following protocol NBA_9168_v114_revL_15Sep2022. After end-prep and cleanup, 50 fmol of each amplicon was used for adapter ligation. The samples were then pooled equimolarly and cleaned using AMPure XP beads. A total of 24.1 fmol of the final pool was loaded onto a Flongle flow cell (FLO-FLG114) and sequenced on a MinION Mk1B for 24 hours using MinKNOW software version 23.11.4. Allele frequencies of G>A in *rob* were calculated based on the raw sequencing data by RStudio.

### Statistical analysis

Each experiment included technical triplicates, and results were obtained from at least three independent biological replicates (n ≥ 3). Data are presented as mean ± SD. Statistical significance was assessed using two-tailed, unpaired Student’s *t* test. *P* < 0.05 was considered statistically significant. Exact *n* values and statistical tests are indicated in the figure legends.

## Results

### Evolutionary analysis and structure of Rob proteins from *Yersinia enterocolitica*

The *Y. enterocolitica* genome encodes two proteins annotated as Rob, referred to here as Rob_1 and Rob_2 (**Fig. 1A**). Rob_1 is the transcriptional regulator in which we observed a high-frequency upstream mutation (-57 G>A). To better understand the evolutionary context of Rob in *Y. enterocolitica*, we aligned 14 AraC-family protein sequences (Rob, MarA, SoxS, and RamA) across *Enterobacteriaceae* species. Phylogenetic analysis revealed that Rob_2 shares high protein identity with typical Rob homologs found in *Y. pestis*, *Y. pseudotuberculosis*, *K. pneumoniae*, *E. cloacae*, *S. enterica*, *C. freundii*, *E. coli*, and *S. flexneri* (**Fig. 1B–D**). In contrast, Rob_1 exhibits only moderate N-terminal identity with other AraC family proteins, such as MarA, SoxS, and RamA. Since *rob_1* is conserved across 94 *Y. enterocolitica* strains (**Fig. S1**), we conclude that Rob_1 and Rob_2 represent two distinct AraC-family proteins. Rob_2 likely evolved from *Enterobacteriaceae* species, whereas Rob_1 appears to be a lineage-specific conserved protein in *Y. enterocolitica*.

**Fig. 1.**
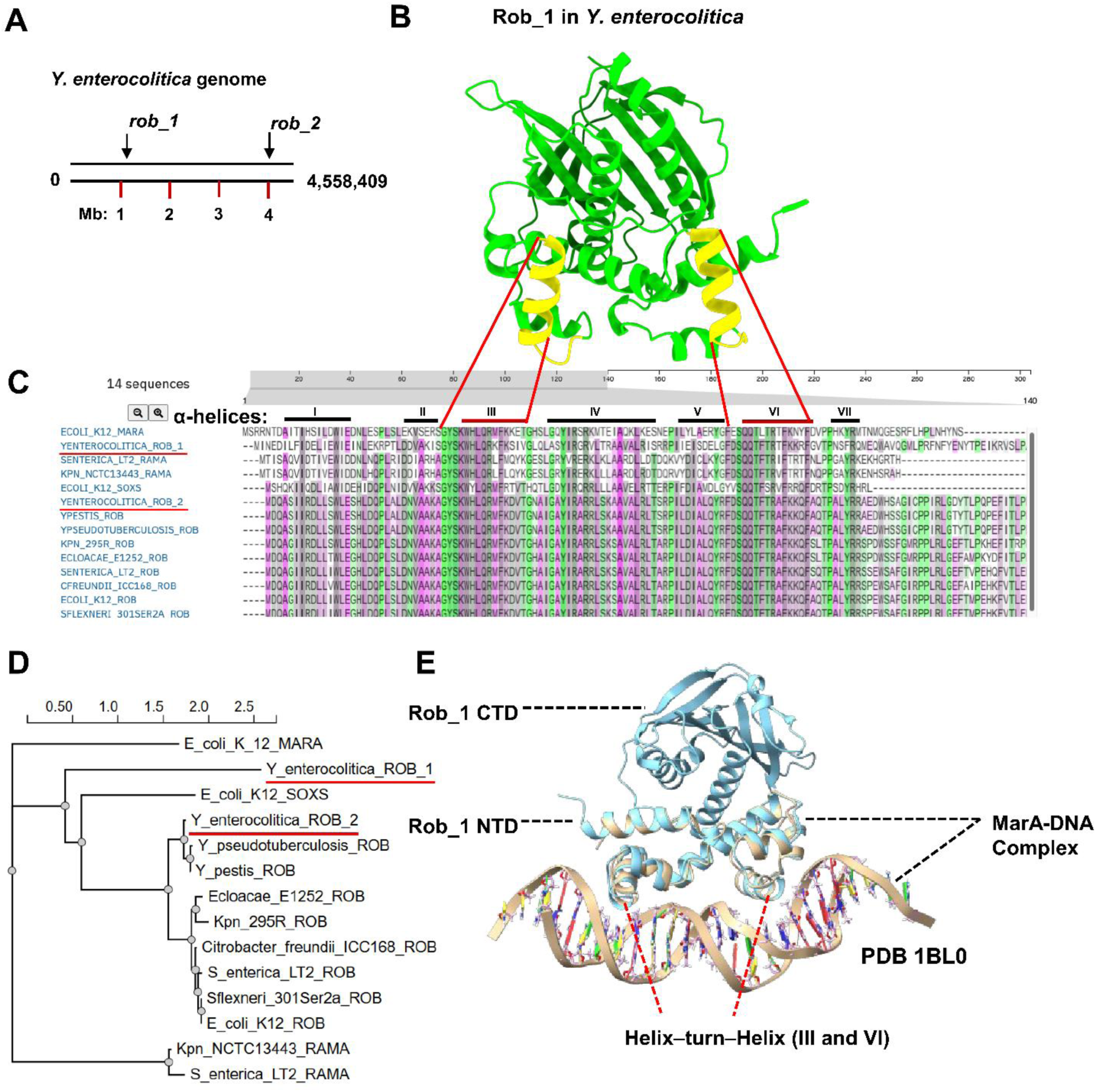
Comparative analysis and structural characterization of Rob_1 in *Yersinia enterocolitica*. **(A)** Genomic locations of *rob_1* and *rob_2* on the *Y. enterocolitica* chromosome. **(B)** Predicted 3D structure of Rob_1 by AlphaFold2 **(C)** Multiple sequence alignment of Rob_1 with homologous AraC-family regulators (MarA, RamA, SoxS, and Rob) from 14 representative Enterobacteriaceae species, including *E. coli* K12 (MarA, SoxS, Rob), *Y. enterocolitica* (Rob_1, Rob_2), *Y. pestis* (Rob), *Y. pseudotuberculosis* (Rob), *K. pneumoniae* NCTC13443 (RamA), *S. enterica* LT2 (RamA, Rob), *S. marcescens* Db11 (Rob), *E. cloacae* E1252 (Rob), *C. freundii* ICC168 (Rob), and *S. flexneri* 301Ser2a (Rob). Conserved domains III and VI, corresponding to DNA-binding helices within the N-terminal domain, exhibit high sequence conservation across species. **(D)** Phylogenetic tree showing the evolutionary relationships of Rob_1 and Rob_2 with related AraC-family regulators. **(E)** Structural model of Rob_1 bound to DNA. Rob_1 is mapped onto the first resolved MarA–DNA complex using the conserved α-helices III and VI. The model highlights the helix–turn–helix (HTH) motif within the N-terminal domain and the predicted DNA-contacting residues inferred from the MarA–DNA structure.

To explore the DNA binding by Rob_1 in *Y. enterocolitica*, we aligned the six DNA-binding domains of MarA with those of the 14 AraC-family. The α-helices III and VI were highly conserved among all AraC-family proteins across species. These two domains form helix–turn–helix (HTH) motifs that insert into adjacent major groove segments of DNA to initiate transcription. To investigate the regulatory mechanism of Rob_1 in *Y. enterocolitica*, we aligned the AlphaFold2-predicted structure of Rob_1 with the resolved *E. coli* MarA–DNA complex structure. The analysis showed that the DNA-binding domain of *Y. enterocolitica* Rob_1 (N-terminal) is highly congruent with that of *E. coli* MarA (115 of 129 residues aligned; RMSD, 0.95 Å; TM-score, 0.94) (**Fig. 1E**). Consequently, Rob_1 is referred to simply as "Rob" throughout the rest of this study.

### A high-frequency mutation in *rob* associated with antibiotic resistance evolution

To develop high-level *de novo* antimicrobial resistance, *Y. enterocolitica* was exposed to step-wise increasing concentrations of six antibiotics(30). WGS analysis showed that 6 out of 12 *Y. enterocolitica* strains consistently acquired the same -57 G>A mutation in *rob* promoter during evolution against enrofloxacin, tetracycline, and chloramphenicol. To explore the role of the above-mentioned mutation in *rob* during these three resistance evolutions, the dynamics of the mutation in *rob* upon exposure to the bactericidal antibiotic enrofloxacin and the bacteriostatic compound tetracycline was investigated using PCR and Sanger sequencing. The mutation in *rob* was detected on passages 13 and 17 (out of 26) during enrofloxacin resistance evolution in the two replicates (**Fig. 2A**). However, during tetracycline resistance evolution, it appeared earlier, on passages 5 and 7 (out of 21) (**Fig. 2B)**.

**Fig. 2.**
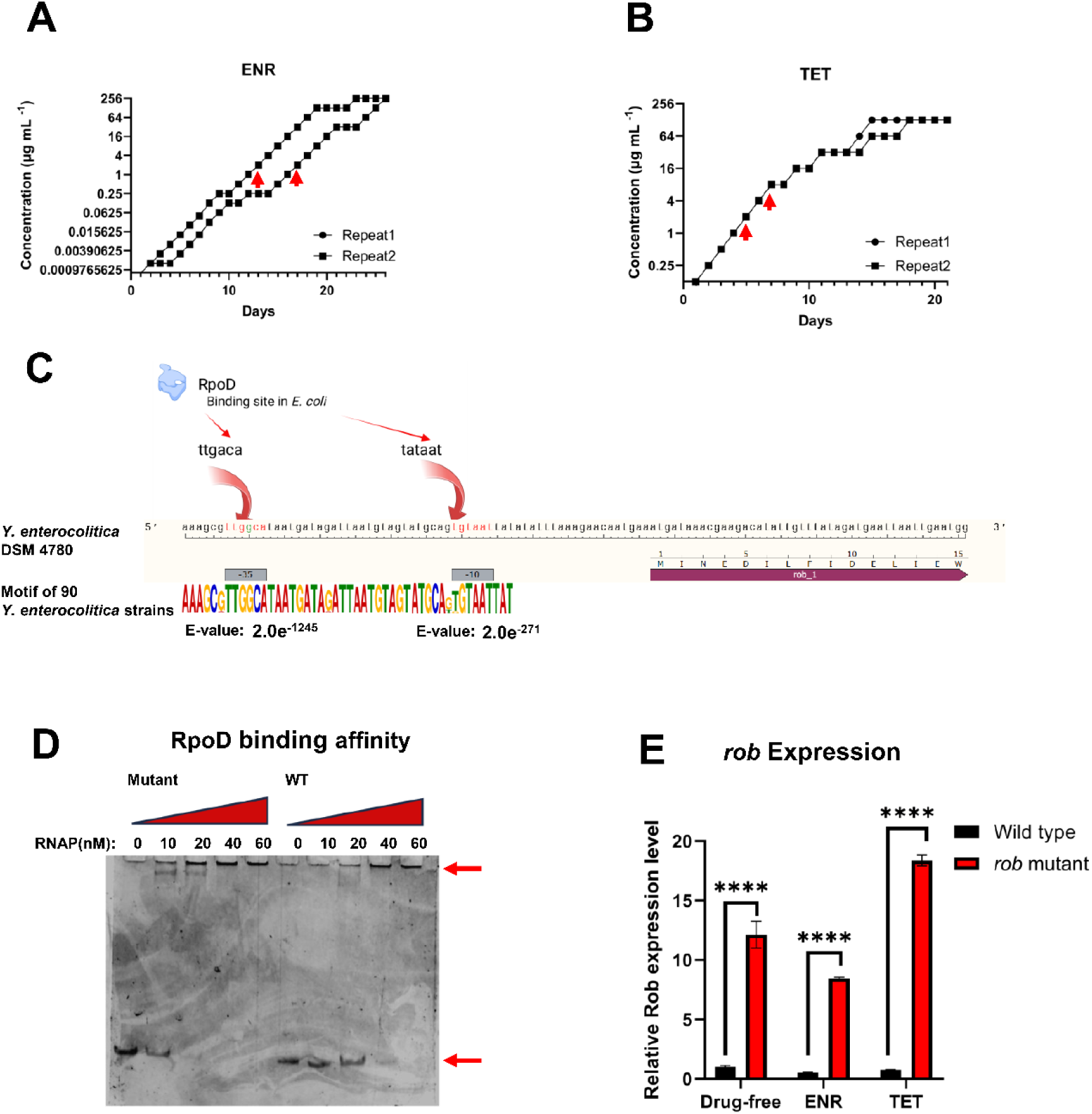
A promoter SNP enhances *rob* expression by increased RpoD binding affinity. A conserved mutation in *rob* -57 G>A acquired during resistance evolution of **A)** enrofloxacin **B)** tetracycline in *Y. enterocolitica.* The day that the promoter mutation in *rob* appeared is indicated by arrows. **C)** Schematic representation of RpoD binding the promoter sequence at the -35 and -10 canonical sequences in *rob*. **D)** Binding affinity test of 150 bp *rob* promoter sequences from *rob* mutant or wild-type strains with RNAP–σ70 holoenzyme (0–60 nM) in EMSA experiment. **E)** Expression levels of *rob* during the stationary phase in the wild-type and *rob* mutant strains in *Y. enterocolitica* under the conditions drug-free, 1/16 MIC of enrofloxacin, and 1/8 MIC of tetracycline, **** P < 0.0001, unpaired two-tailed Student’s *t*-test.

### A promoter SNP enhances *rob_1* expression by increasing RpoD binding affinity

The expression of *rob* in *E. coli* is regulated by RpoD or RpoS in response to environmental stimuli(32, 33). Aligning *E. coli* RpoD binding sites with the *rob* promoter sequence in *Y. enterocolitica* revealed that the mutation is located within the -35 canonical sequence (TTGACA), creating a perfect match following the -57 G>A transition (**Fig. 2C**). The canonical *rob* promoter sequence is highly conserved across *Yersinia enterocolitica* strains (**Fig. 2C**). To determine whether the promoter mutation in *rob* of *Y. enterocolitica* impacts the RNA polymerase binding affinity, we respectively incubated RpoS and RpoD proteins with the 150 bp *rob* promoter sequence from the wild-type and *rob* mutant (-57 G>A) strains in an Electrophoretic Mobility Shift Assay (EMSA). No clear shift was observed when RpoS was incubated with the *rob* promoter sequences either from the wild-type strain or the mutant strain (**Fig. S2**). However, the mutant *rob* promoter sequence bound to RpoD at 20 nM in the gel shift analysis, exhibiting a 2- to 3-fold higher binding affinity compared to the wild-type strain (**Fig. 2D**).

To quantify the effect of the single -57 G>A mutation in *rob* in *Y. enterocolitica*, we constructed a strain carrying only the *rob* -57 G>A mutation. qPCR analysis in the stationary phase showed that the mutation enhanced *rob* expression under drug-free conditions and during exposure to subinhibitory concentrations of enrofloxacin and tetracycline (*p* < 0.0001) (**Fig. 2E**). MIC analysis of ten antibiotic classes showed that the *rob* promoter mutation increased resistance 4-to 8-fold for five antibiotics, while decreasing it 2- to 8-fold for the remaining five (**Table 1)**.

**Table 1.** Effect of *rob* overexpression on susceptibility to ten commonly used antibiotics.

| Antibiotic | MIC WT<br>($\mu\text{g/mL}$) | MIC *rob* mutant<br>($\mu\text{g/mL}$) | Fold change<br>(mutant/WT) |
| --- | --- | --- | --- |
| Enrofloxacin | 0.0625 | 0.25 | 4× |
| Tetracycline | 1 | 8 | 8× |
| Amoxicillin | 32 | 128 | 4× |
| Erythromycin | 128 | 512 | 4× |
| Chloramphenicol | 4 | 32 | 8× |
| Cefepime | 4 | 2 | 0.5× |
| Cefotaxime | 8 | 2 | 0.25× |
| Fosfomycin | 1024 | 128 | 0.125× |
| Rifampicin | 256 | 64 | 0.25× |
| Kanamycin | 64 | 32 | 0.5× |

### Rob directly binds promoters to activate efflux-related multidrug resistance genes

To identify the genome-wide binding sites of Rob in *Y. enterocolitica*, we constructed a C-terminal FLAG tag at the native *rob* locus, which was confirmed by Western blotting (**Fig. S3**). ChIP-seq was performed to map the genome-wide binding of Rob-3F during the early exponential phase of cells growing at 1/8 MIC of tetracycline. An untagged wild-type strain grown under the same conditions served as a control. Peak calling using MACS2 identified four strong peaks (*meoA*, *tolC*, *mlaF*, and *atpI*) (**Fig. 3A-D**) and four weak peaks (*limB*, *NCTC12982_01318*, *lpxC*, and *ydhP_3*). *meoA*, *tolC*, *mlaF*, and *ydhP_3* are associated with outer-membrane homeostasis and efflux activity; *atpI* encodes an *F_0_ F_1_*-ATP synthase assembly factor; while *limB*, *NCTC12982_01318*, and *lpxC* are involved in buffering oxidative and metal stress. MEME analysis of these eight peaks identified a weakly conserved palindromic motif (**Fig. S4**).

**Fig. 3.**
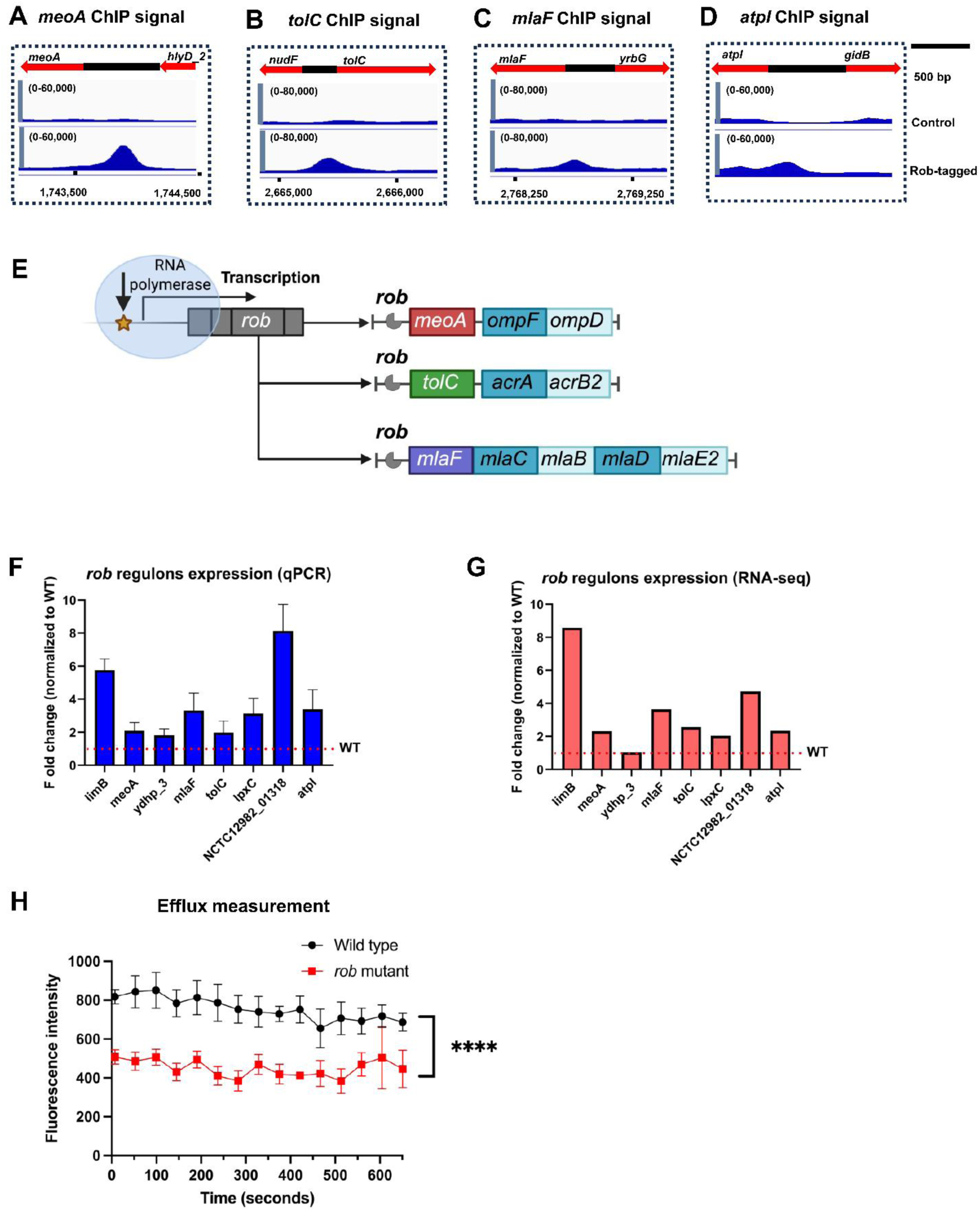
Identification of Rob binding sites in *Y. enterocolitica* by ChIP-seq. **(A–D)** ChIP-seq profiles showing Rob binding at the promoter regions of *meoA* (A), *tolC* (B), *mlaF* (C), and *atpI* (D). Blue tracks represent normalized ChIP-seq signal in the control strain (top) and Rob-tagged strain (bottom). Gene orientations are indicated by arrows; scale bar, 500 bp. **(E)** Schematic model of Rob-dependent transcriptional regulation. Rob binding upstream of target operons activates genes involved in outer membrane permeability (*ompF/ompD*), multidrug efflux (*tolC–acrAB2*), and phospholipid transport (*mla* operon). **(F)** Relative expression of selected Rob regulon genes measured by qPCR, shown as fold change normalized to wild type (WT). Data represent mean ± SD. The red dashed line indicates WT expression level. **(G)** Corresponding fold changes in gene expression derived from RNA-seq analysis, normalized to WT, confirming global upregulation of Rob target genes. **(H)** Efflux activity measured over time using a fluorescence-based assay. Rob mutant exhibits significantly higher efflux compared to the WT. Data are shown as mean ± SD; **** indicates *P* < 0.0001.

The *Y. enterocolitica* DSM 4780 *rob* -57 G>A strain, which exhibits increased *rob* expression compared with the parental strain and is therefore designated the *rob* overexpression strain, was used to assess Rob-dependent transcriptional regulation. Using qPCR, we found that the expression of eight candidate Rob-regulated genes: *meoA, tolC, mlaF, atpI, limB, NCTC12982_01318, lpxC,* and *ydhP_3* was higher in the *rob* mutant than in the wild-type strain during exponential growth (**Fig. 3F**). This expression pattern was independently confirmed by RNA-seq analysis. Together, these data support a role for Rob as a transcriptional activator of these genes (**Fig. 3G**).

Given the identification of Rob binding and the elevated efflux gene expression observed in the *rob* mutant, efflux pump activity was assessed using Nile Red dye as a fluorescent substrate in a time-dependent efflux assay. The *rob* mutant strain exhibited significantly lower fluorescence retention (*p* < 0.0001) for Nile Red compared to the wild-type strain (**Fig. 3H, S5**). In conclusion, *rob* is an efflux transcriptional regulator in *Y. enterocolitica*.

### *rob* overexpression upregulates Fe–S cluster and heme biosynthesis

RNA-seq was used to profile transcriptional changes associated with *rob* overexpression during efflux activation (**Fig. S6**). Under 1/16 MIC enrofloxacin, *rob* expression increased 10.6-fold and 260 genes were upregulated, spanning transcription, signal transduction, membrane biogenesis, carbohydrate metabolism, and inorganic-ion transport (**Fig. 4A**). Pathways supporting ATP production via the respiratory chain were strongly induced, including Fe–S cluster homeostasis and heme biosynthesis. Heme biosynthesis genes, *hemB* (1.8-fold) and *hemN_3* (26.2-fold); heme uptake/transport genes, *hemR* (5.3-fold), *hemP* (9.9-fold), *hemS* (2.1-fold), NCTC12982_03534 (39.1-fold); redox genes, *qor2* (6.9-fold), NCTC12982_03533 (19.8-fold), *nirB* (5.1-fold), *nirC* (6.3-fold), *nirD* (13.9-fold) and ferric uptake/transport genes, *ttrA* (7.1-fold), *ttrB* (78.8-fold), *ttrC* (20.5-fold), *fhuF* (5.0-fold) were all significantly upregulated (**Fig. 4C**; **Table 2**). Conversely, 214 genes related to translation and amino-acid metabolism were downregulated (**Fig. 4B**).

**Fig. 4.**
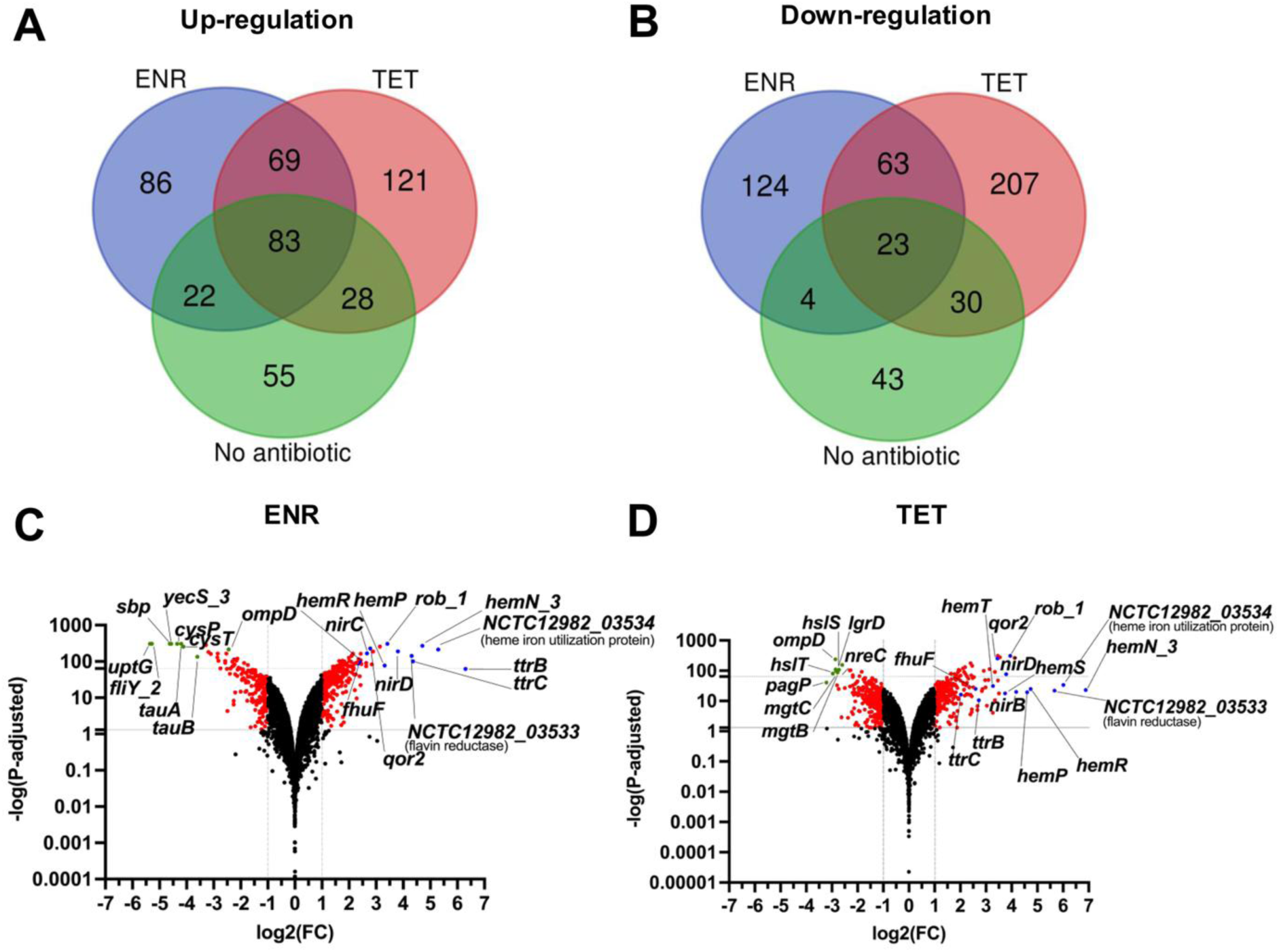
Transcriptomic analysis of expression changes triggered by overexpression of *rob*. **A)** Expression of upregulated genes quantified by RNA-seq in the *rob* mutant compared to wild-type under drug-free, 1/16 MIC enrofloxacin, and 1/8 MIC tetracycline. **B)** Expression of downregulated genes quantified by RNA-seq in the *rob* mutant compared to wild-type under drug-free conditions, 1/16 MIC enrofloxacin, and 1/8 MIC tetracycline. **C)** The volcano plot of up and downregulation of *rob* mutant strain compared to wild-type strain exposed to ENR. **D)** The volcano plot of up and downregulation of *rob* mutant strain compared to wild-type strain exposed to TET.

**Table 2.** The upregulated genes involved in Fe–S cluster homeostasis and heme biosynthesis.

| Reactions | Gene | ENR (FC) | TET (FC) | Function |
| --- | --- | --- | --- | --- |
| **Heme biosynthesis** | *hemB* | 1.8 | 2.7 | Delta-aminolevulinic acid dehydratase |
|  | *hemN_3* | 26.2 | 117.8 | Coproporphyrinogen-III oxidase |
| **Heme intake and transport** | *hemP* | 9.9 | 24.3 | Hemin uptake protein |
|  | *hemR* | 5.3 | 26.7 | Hemin receptor |
|  | *hemU* | 1.4 | 7.1 | Hemin transport system permease |
|  | *hemT* | 1.7 | 9.7 | Hemin-binding periplasmic protein |
|  | *hemS* | 2.1 | 13.4 | Hemin transport protein |
|  | *NCTC12982_03534* | 39.1 | 50.6 | Heme iron utilization protein |
| **Oxidation-reduction reaction** | *qor2* | 6.9 | 10.9 | Quinone oxidoreductase |
|  | *NCTC12982_03533* | 19.8 | 65.5 | Flavin reductase |
|  | *sodA* | 3.3 | 5.2 | Superoxide dismutase |
|  | *nirB* | 5.1 | 18.0 | Nitrite reductase |
|  | *nirC* | 6.3 | 7.9 | Nitrite transporter |
|  | *nirD* | 13.9 | 13.7 | Nitrite reductase small subunit |
| **Tetrathionate reductase** | *ttrA* | 7.1 | 2.1 | Tetrathionate reductase subunit TtrA |
|  | *ttrB* | 78.8 | 6.5 | Tetrathionate reductase subunit B |
|  | *ttrC* | 20.5 | 4.1 | Tetrathionate reductase subunit TtrC |
| **Ferric enterobactin transport** | *fepB* | 1.8 | 2.6 | Ferric enterobactin-binding periplasmic protein |
|  | *fepC* | 2.5 | 2.9 | Ferric enterobactin transport ATP-binding protein |
|  | *fepD* | 2.6 | 2.6 | Iron-enterobactin transporter membrane protein |
|  | *fepG* | 2.5 | 2.4 | Ferric enterobactin ABC transporter, permease protein |
| **Ferric hydroxamate uptake** | *fhuA* | 1.0 | 3.6 | Hydroxamate-type ferrisiderophore receptor |
|  | *fhuC_1* | 2.9 | 2.8 | ABC transporter ATP-binding protein |
|  | *fhuC_2* | 1.8 | 2.1 | ABC transporter ATP-binding protein |
|  | *fhuD* | 2.2 | 2.1 | Iron-hydroxamate transporter substrate-binding subunit |
|  | *fhuF* | 5.0 | 6.0 | Siderophore-iron reductase FhuF |

Under 1/8 MIC of tetracycline, *rob* increased 15.4-fold,upregulating 301 genes (**Fig. 4A**). The induction of ATP-producing, heme-associated redox pathways was even more pronounced than under enrofloxacin conditions: heme biosynthesis (*hemB*, 2.7-fold; *hemN_3*, 118-fold), heme uptake/transport (*hemR*, 26.7-fold; *hemP*, 24.3-fold; *hemS*, 13.4-fold; NCTC12982_03534, 50.6-fold), redox genes (*qor2*, 10.9-fold; NCTC12982_03533, 64.5-fold; *nirB*, 18.0-fold; *nirC*, 7.9-fold; *nirD*, 13.7-fold), and ferric uptake/transport (*ttrB*, 6.5-fold; *ttrC*, 4.1-fold; *fhuF*, 6.0-fold) were all significantly upregulated (**Fig. 4D**; **Table 2**). Meanwhile, 323 genes involved in translation, transcription, and inorganic-ion transport were downregulated (**Fig. 4B**). Across conditions, 152 upregulated and 86 downregulated genes overlapped, with correlated effect sizes (R² = 0.439; **Fig. S7**).

### *rob* mutation drives antimicrobial resistance acquisition

To investigate whether the -57 G>A mutation influences resistance acquisition rates, we exposed wild-type (basal expression) and *rob* mutant (overexpression) strains to step-wise increasing concentrations of tetracycline and enrofloxacin.

Upon enrofloxacin exposure, the *rob* -57 G>A mutation altered the evolutionary trajectory, specifically the order of mutation accumulation, rather than the final resistance level. In the wild type, resistance increased gradually, catching up to the *rob* mutant’s MIC by day 9 (**Fig. 5A**). WGS of populations from days 3–20 revealed that resistance was driven by mutations in *rob*, *gyrA*, *gyrB*, and *parC* in both backgrounds. By day 9, both strains carried the *gyrA* S83I and *rob* -57 G>A genotype and exhibited the same MIC. However, due to the step-wise protocol initiating at different absolute concentrations, the wild-type was exposed to an antibiotic concentration eight-fold lower than the *rob* mutant at this time point (**Fig. S8B**). Under lower concentration pressure, the *rob* mutation was transiently lost in wild-type lineages. With step-wise increases in enrofloxacin exposure, the *rob* mutation fixed again in the final wild-type lineages. In mutant lineages evolved under constant enrofloxacin exposure, the mutation was never lost (**Fig. 5B**).

**Fig. 5.**
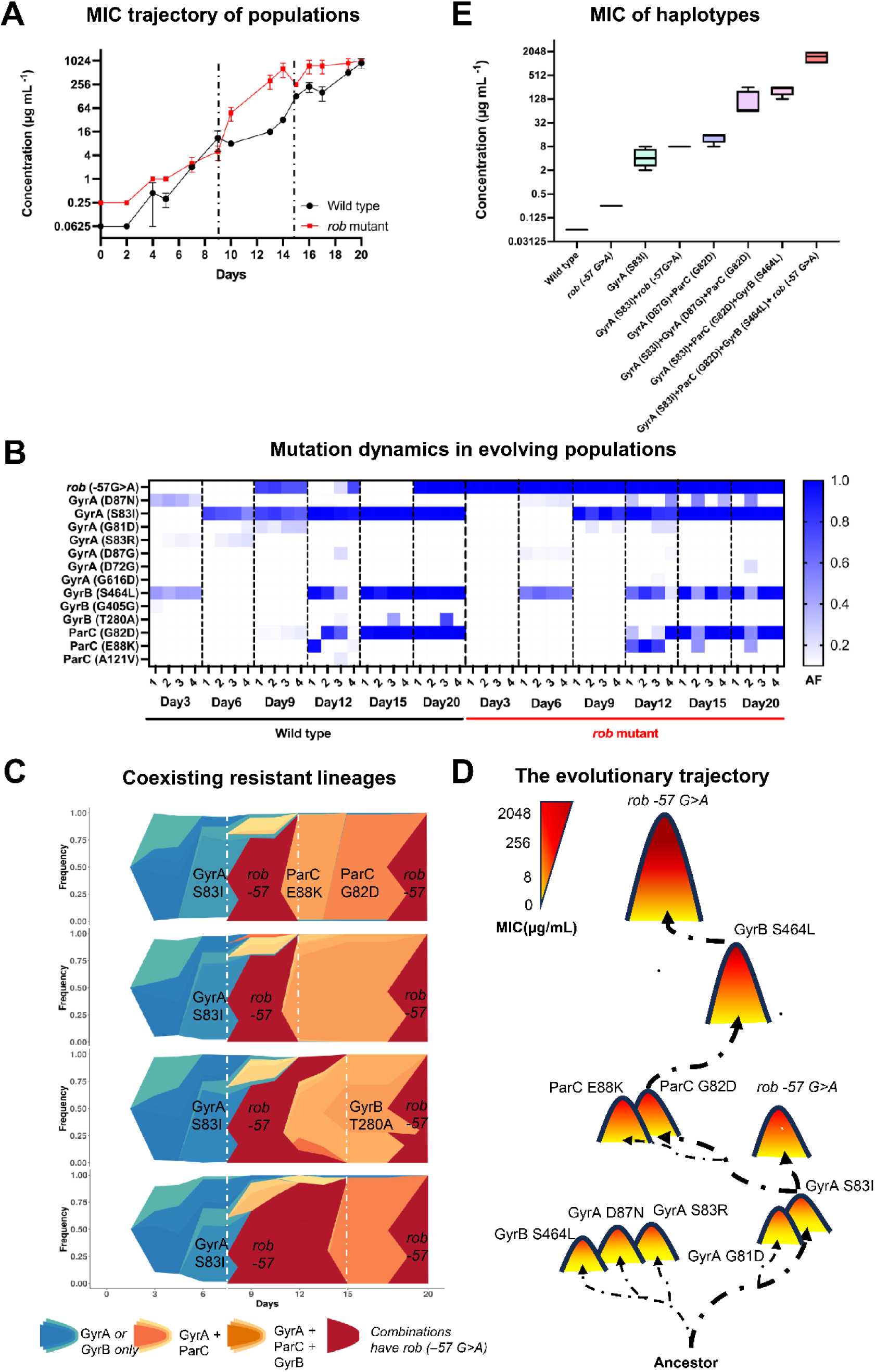
Enrofloxacin resistance acquisition in wild-type and *rob* mutant strains. **A)** Enrofloxacin MIC trajectories for evolving populations. *rob* mutant lineages altered the trajectory of resistance acquisition. Dashed lines indicate time points of clonal isolation. **B)** Allele-frequency heat map over 20 days of evolution. The *rob* -57 G>A mutation arises early, followed by mutations in GyrA, ParC, and GyrB. Wild-type and *rob* mutant founders are separated by a dashed line. **C)** Muller plots of wild-type populations showing the rise of resistant lineages. Blue: GyrA or GyrB only; green: GyrA and ParC; orange: GyrA, ParC and GyrB; red: any genotype carrying *rob* -57 G>A. White dashed lines mark days 9 and 15. **D)** Schematic of the evolutionary trajectory toward high-level resistance: early acquisition of *rob* -57 G>A followed by sequential mutations in target genes (GyrA S83I, ParC G82D, GyrB S464L). **E)** MICs of representative haplotypes, illustrating step-wise resistance increases with the addition of *rob*, then ParC or GyrB mutations. Red dashed lines separate genotypic groups.

To analyse these dynamics, we reconstructed lineage competition using Muller plots derived from population sequencing and colony isolation. Early evolution was characterized by a soft selective sweep of *gyrB* and *gyrA* variants, with *gyrA* S83I becoming dominant. The *rob* -57 G>A mutation initially arose on this *gyrA* S83I background. However, under intermediate selection (days 9–15), lineages carrying combinations of *gyrA* and *parC*, or *gyrA*, *gyrB*, and *parC*, outcompeted the *gyrA* S83I + *rob* lineage (**Fig. 5C-D**). Colony isolation confirmed that while *gyrA* S83I + *rob* dominated on days 10–11 (**Table 3**), it was displaced by *parC*-containing genotypes between days 12 and 15, which conferred higher resistance and superior growth under low enrofloxacin concentrations (**Fig. 5E and Fig. S9**). Ultimately, *rob* -57 G>A was reacquired and fixed on the high-fitness *gyrA* S83I + *gyrB* S464L + *parC* G82D background in the final lineages. Thus, the *rob* -57 G>A lineage was transiently competitive but displaced during intermediate evolution, only to be re-selected as essential for tolerating maximal enrofloxacin exposure.

**Table 3.** Genotypes from 40 colony isolations.

| Haplotype | Isolates |
| --- | --- |
| Wild type | Wild type |
| GyrA (S83I) | 10,16,17,20 |
| GyrA (S83I) + *rob* -57 G>A | 3-5,9,26-28,30-32,19,22,24,25,34,37-40 |
| GyrA (D87G) + ParC (G82A) | 2,21,29,33,35,36 |
| GyrA (S83I) + GyrA (D87G) + ParC (G82D) | 6,12,13,14,15 |
| GyrA (S83I) + GyrB (S464L) + ParC (G82D) | 7,8 |
| GyrA (S83I) + GyrB (S464L) + ParC (G82D) + *rob* -57 G>A | Final population isolates |

Under step-wise tetracycline exposure, the wild-type strain rapidly reached the resistance level of the *rob* mutant by day 7, with converging evolutionary trajectories (**Fig. S10A**). Sequencing detected no gene amplifications but identified mutations in *rob*, *bcr*, *acrB*, *acrR*, *ompD*, and *nusE*. The *rob* -57 G>A mutation emerged within four days in the wild type and fixed in most replicates. Notably, *rob* overexpression biased evolution toward efflux mechanisms: mutant populations acquired more mutations in efflux genes (*bcr*, *acrB*, and *acrR*) and fewer in the ribosome-associated target gene *nusE* (V57L) compared to the wild type (**Fig. S10B**). These patterns indicate that *rob* overexpression biases tetracycline resistance evolution toward efflux-based mechanisms rather than target modification.

### Rapid counter-selection of the *rob* -57 G>A mutation under drug-free conditions

*rob* overexpression imposed a measurable fitness cost (approximately 9.6% reduction in growth rate) under drug-free conditions (**Fig. S11**), although it conferred a growth advantage in the presence of 1/16 MIC enrofloxacin or 1/8 MIC tetracycline (**Fig. 6A**). To assess stability, we propagated the mutant strain without antibiotics for 15 days. Enrofloxacin and tetracycline resistance rapidly declined to wild-type levels during this period (**Fig. 6B-C**). Amplicon sequencing showed that the frequency of the *rob* −57 G>A allele decreased, whereas the wild-type allele increased within seven days (**Fig. 6D**).

**Fig. 6.**
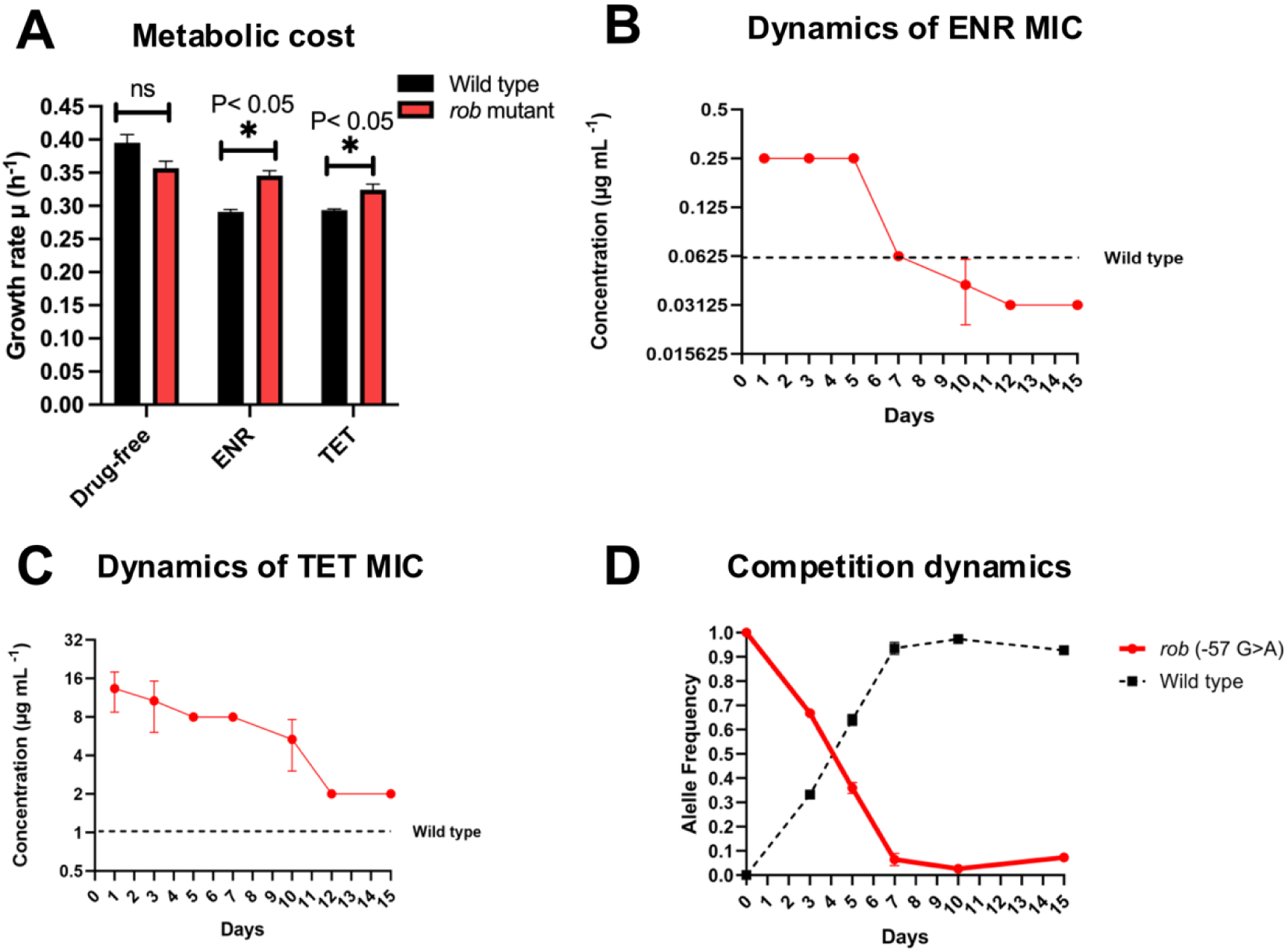
Loss of the *rob* -57 G>A mutation during 15 days of drug-free growth. **A)** The fitness cost in the form of reduced growth rate due to *rob* overexpression measured in three conditions. **B)** Enrofloxacin MIC of *rob* mutant during drug-free adaptation. **C)** Tetracycline MIC of the *rob* mutant during drug-free adaptation. **D)** Allele frequency of the mutation -57 G>A in *rob* for 15 days adaptation.

To determine whether the decrease in the *rob* −57 G>A allele upon drug withdrawal resulted from genetic back-mutation or population-level counter-selection, we sequenced 14 individual colonies isolated between days 3 and 5 of drug-free passage. Colonies that lost the phenotype shared an identical wild-type genomic background, whereas resistant colonies retained the −57 G>A mutation and the original mutant background. These results confirm that the rapid loss of the variant resulted from counter-selection against the costly mutant genotype rather than genetic reversion (**Table S2**).

## Discussion

The combined outcome of this study characterizes a novel resistance mechanism in *Yersinia enterocolitica*, driven by a promoter SNP in the transcriptional regulator *rob*. Despite the low spontaneous mutation rate (∼10⁻⁹ bp⁻¹ per cell per generation)(34, 35), this regulatory mutation functions as an early adaptive hub that biases subsequent evolutionary trajectories under antibiotic selection. Integrated genomic, transcriptomic, and EMSA data show that the mutation increases RpoD binding, driving strong *rob* overexpression. Elevated Rob activates downstream efflux systems across multiple regulons, broadening the substrate export capacity. By constructing both ‘basal’ and ‘overexpression’ models of the *rob* regulatory network, we demonstrate that modulation of a single transcription factor, Rob, can accelerate pathogen evolvability under antibiotic stress.

Rob in *Y. enterocolitica* is an AraC/XylS-family transcription factor that has not previously been functionally defined in this species. Sequence and structural comparisons reveal high N-terminal conservation with AraC/XylS homologs, consistent with efflux-associated regulatory functions reported in related bacteria(16). We confirmed that *rob* overexpression increased efflux efficiency in *Y. enterocolitica*. The divergence in the C-terminal alignment may contribute to differences in DNA-binding specificity. In *E. coli*, Rob pre-recruits RNA polymerase to the *marA-soxS-rob* binding site in its promoter region to initiate transcription(23), regulating cellular responses to superoxide, organic solvents, and heavy metals(36). However, no such binding site was examined in the *rob* promoter of *Y. enterocolitica*. EMSA experiments revealed that RpoD exhibited stronger binding affinity to the mutant allele (TTGACA) at the -35 box of the *rob* promoter, suggesting *rob* transcription in *Y. enterocolitica* is primarily RpoD-dependent. ChIP-seq identified a limited set of Rob-associated binding sites under the tested conditions. Thus, unlike in *E. coli* where Rob functions more broadly(36), Rob in *Y. enterocolitica* has evolved to serve primarily as an efflux-focused regulator during long-term evolution.

While copy number variation (CNV) is a common driver of resistance(37–39), transcription factor-mediated overexpression engages similarly complex regulatory mechanisms. Rob acts as a central regulatory node coordinating multiple efflux- and membrane-associated pathways. Consequently, *rob* overexpression triggers a cascade of efflux- and influx-related responses. Transcriptomic analysis suggests energetic compensation for increased efflux, with Fe–S cluster and heme biosynthesis upregulated and ribosomal protein expression reduced (**Fig. 7**). Overall, CNV-driven and regulator-driven mechanisms produced comparable increases in efflux capacity. Both mechanisms can upregulate efflux pathways and increase resistance. However, the breadth of cross-resistance depends on the substrate range of the induced efflux systems(40). Overexpression of the narrow-spectrum efflux pump NorA does not confer noticeable cross-resistance to other classes of antibiotics(25). In contrast, the *rob* mutation upregulates efflux pumps from MLA, OMP, and RND families, which have broad substrate specificities. Accordingly, we observed cross-resistance to 5 of 10 antibiotics tested, supporting the conclusion that the *rob* variant acts as a multidrug transcriptional regulator.

**Fig. 7.**
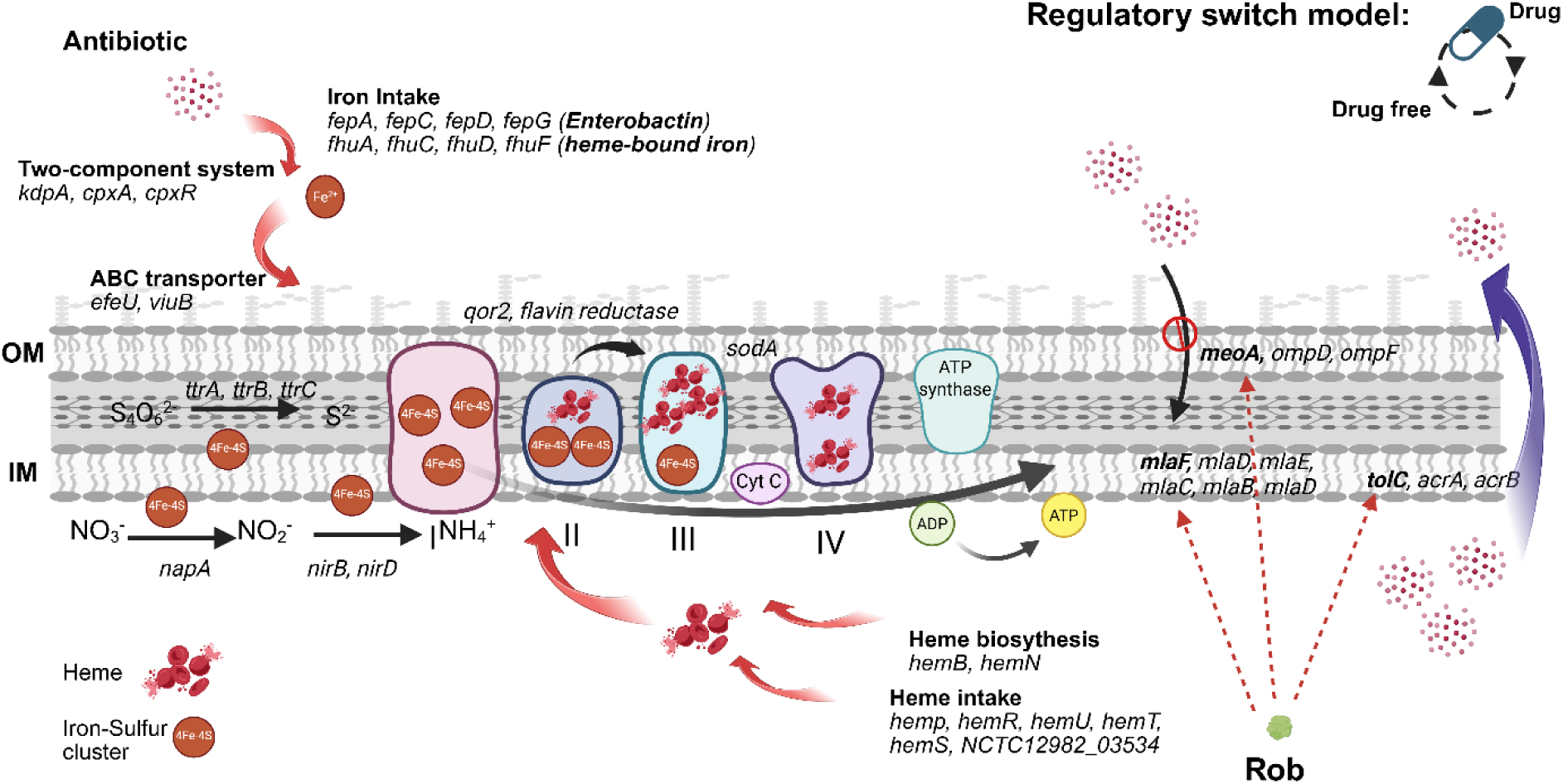
Mechanism: *rob* overexpression activated Fe–S cluster homeostasis and heme biosynthesis. In *Y. enterocolitica*, overexpression of *rob* induces transcriptional activation of genes involved in iron acquisition (*fep*, *fhu*, *viuB*), iron-sulfur (4Fe–4S) cluster assembly (*napA*, *nirB*, *ttrABC*), heme biosynthesis (*hemB*, *hemN*), and heme uptake (*hemP*, *hemR*, *hemU*, *hemT*, *hemS*). These regulatory changes enhance respiratory chain function (complexes I–IV), ATP production, and the expression of efflux systems (*acrAB, tolC*, *ompD*, *mlaFEDB*) in response to antibiotic exposure. The regulatory switch is driven by clonal competition: drug exposure enriches resistant, high-cost clones, whereas drug-free growth counter-selects them in favor of sensitive, low-cost clones.

Interactions between regulatory and structural loci can profoundly shape evolutionary trajectories during antibiotic adaptation(41–43)(26). While transcriptional regulators allow cells to fine-tune responses to environmental stimuli(44, 45), our data suggest that the interplay between *rob* and downstream targets differs depending on the antibiotic context, driving distinct evolutionary outcomes.

In tetracycline resistance evolution, the early mutation in *rob* acts as an adaptive ‘hub’ that reshapes the subsequent mutational landscape. Rather than strictly constraining the path, *rob* overexpression biased the trajectory toward efflux-mediated mechanisms. The introduction of the -57 G>A *rob* mutation modified the final mutational profile in *Y. enterocolitica*: by rendering the target modification NusE (V57L) less necessary, the *rob* mutant lineage accumulated a higher frequency of mutations in efflux pump genes (*acrR* and *acrB*) compared to the wild-type. This shift suggests that *rob* overexpression relieves the selection pressure on ribosomal targets, effectively steering the population toward an efflux-centric resistance strategy.

The dynamics of the *rob* -57 G>A mutation during enrofloxacin evolution highlight a complex trade-off between resistance capability and metabolic cost, rather than simple positive epistasis. The mutation imposes a ∼9.6% reduction in growth rate under drug-free conditions. Consequently, its evolutionary fate is strictly governed by the intensity of selection. Under weak selection, the lineage carrying *rob* -57 G>A and *gyrA* (S83I) was outcompeted by lineages carrying target-site combinations (e.g., *gyrA* + *parC*) that conferred comparable resistance with a lower metabolic cost. This indicates that the costly *rob* mutation is competitively disadvantaged when lower-cost structural mutations are sufficient. However, as enrofloxacin exposure increased to lethal levels, the *rob* mutation was reacquired and fixed on top of the target-site mutations. This pattern suggests a relationship of functional complementarity: at high drug concentrations, target-site modifications alone are insufficient, and *rob*-mediated efflux becomes a prerequisite for survival(25). Our data indicate that *rob* -57 G>A is recurrently selected not because it synergistically increases growth rates, but because it provides a necessary survival boost that bridges the gap between the limit of target-site resistance and the lethal antibiotic concentration.

Notably, the high fitness cost of *rob* overexpression, manifested as reduced growth rates, drives rapid clonal replacement upon the removal of antibiotic pressure. Although compensatory mutations arose in metabolic genes (*manC*, *coaD*, *mtnK*, and *bioH*), analysis of the genetic backgrounds revealed that the rapid loss of the *rob* mutation was driven by population-level counter-selection, with the wild-type allele reestablished within 50 generations. Considering the evolutionary conservation of the *rob* promoter among *Yersinia* strains, this regulatory mutation likely represents a specialized, species-specific defense mechanism optimized for transient, high-stakes adaptation rather than long-term persistence.

## Data Availability

All sequencing data generated in this study (WGS, RNA-seq, amplicon sequencing, and ChIP-seq) have been deposited in the NCBI Sequence Read Archive (SRA) under BioProject accession number PRJNA1398315.

## Supplementary Data

Supplementary Data are available at NAR Online.

## Author contributions

Xinyu Wang performed the experiments and wrote the manuscript. Taichi Chen contributed to mutant construction and provided suggestions for the manuscript. Martijs Jonker and Wim de Leeuw assisted with sequencing analysis and bioinformatic interpretation. Alphonse de Koster performed bioinformatics and provided suggestions for the manuscript. Gaurav Dugar helped design experiments and provided comments on the manuscript. Benno ter Kuile conceived the overall project and revised the manuscript.

## Acknowledgments

We thank Stanley Brul and Wenxi Qi for their stimulating suggestions. We also thank Selina van Leeuwen for her assistance with DNA sequencing. Students Desi Komen and Sieradj Hendrikson contributed to this project as part of their degree requirements. Xinyu Wang acknowledges the China Scholarship Council for providing a PhD scholarship.

