## Supplementary Material for "Modulation of *rob* expression accelerates development of antibiotic resistance in *Yersinia enterocolitica*"

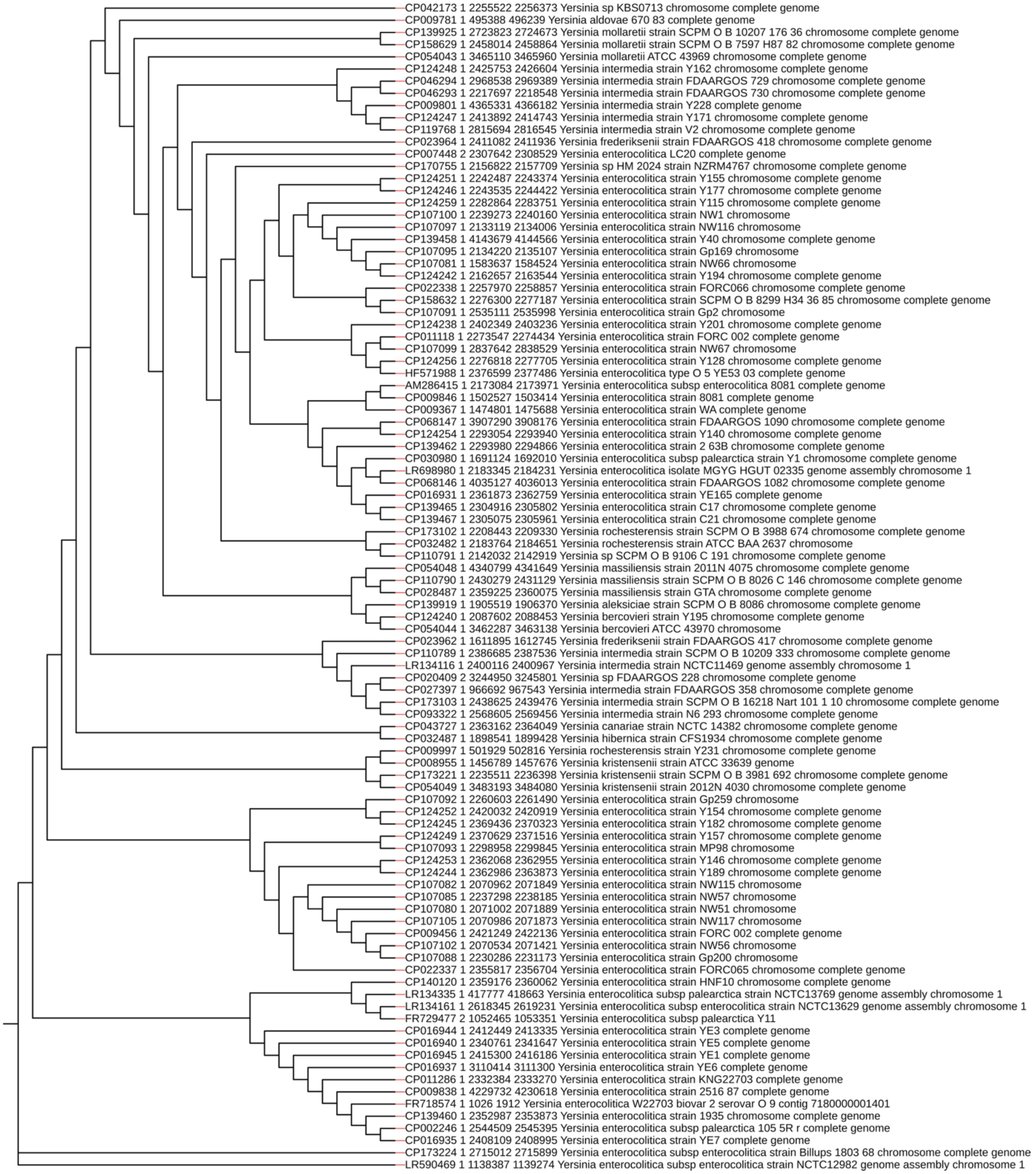


**Fig. S1. Evolutionary analysis of Rob_1 in *Yersinia enterocolitica* based on 94 aligned sequences from NCBI.**

The *rob_1* gene from *Y. enterocolitica* was queried using NCBI Nucleotide BLAST, yielding 94 sequences with identities starting from 79.25%. These aligned sequences were subsequently used for evolutionary analysis with NGPhylogeny.

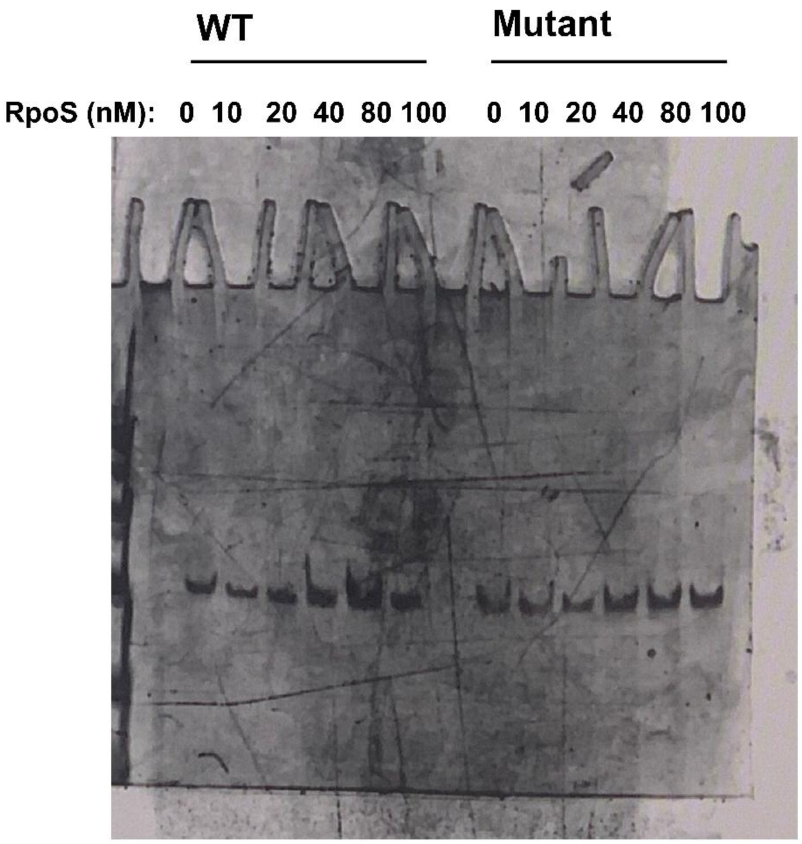


**Fig. S2. RpoS co-incubated with a 150 bp *rob* promoter sequence from wild-type and *rob* mutant strains in EMSA**

Electrophoretic mobility shift assays (EMSA) showing binding of purified RpoS to a 150-bp DNA fragment encompassing the *rob* promoter from the wild-type (WT) and *rob* mutant strains. Increasing concentrations of RpoS (0, 10, 20, 40, 80, and 100 nM) were incubated with the DNA probes prior to separation on a 6% native polyacrylamide gel. The positions of free DNA and RpoS–DNA complexes are indicated by their differential electrophoretic mobilities.

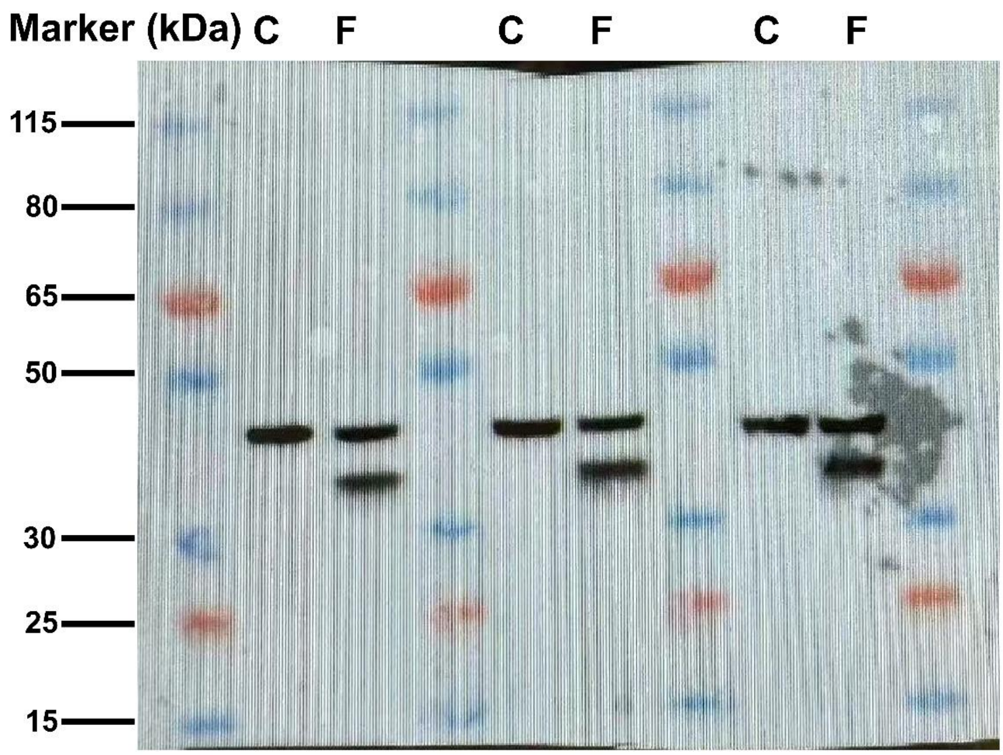


**Fig. S3. Western blot confirmation of the Rob 3×FLAG tagged strain.**

Western blot analysis using an anti-FLAG antibody confirms expression of Rob 3×FLAG tagged strain. Lanes labeled C indicate the control (untagged) strain, while F indicates the Rob 3×FLAG tagged strain. A specific band corresponding to Rob 3×FLAG tagged strain is detected exclusively in the FLAG-tagged samples. Molecular weight markers (kDa) are shown on the left.

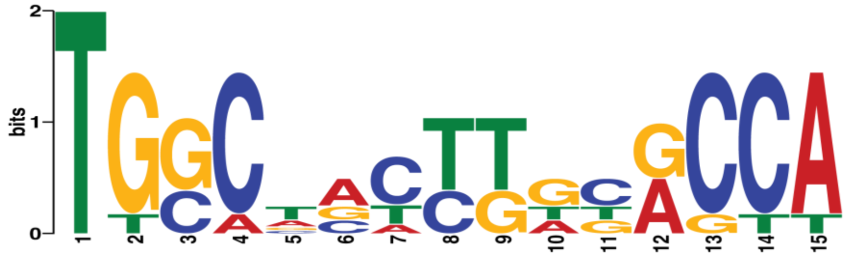


**Fig. S4. ChIP-seq based identification of the Rob DNA-binding motif in *Yersinia enterocolitica* using MEME**

ChIP-seq analysis of Rob binding in *Y. enterocolitica* was performed using MACS2 peak calling. Four high-confidence Rob-binding peaks were detected upstream of *meoA*, *tolC*, *mlaF*, and *atpI* together with four lower-confidence peaks associated with *limB*, *NCTC12982_01318*, *lpxC*, and *ydhP_3*. These eight MACS2 peaks were used as input for *de novo* motif discovery using MEME Suite, identifying the Rob DNA-binding motif. The E-value ($7.9 \times 10^0$) reflects low confidence as only 8 peaks were identified.

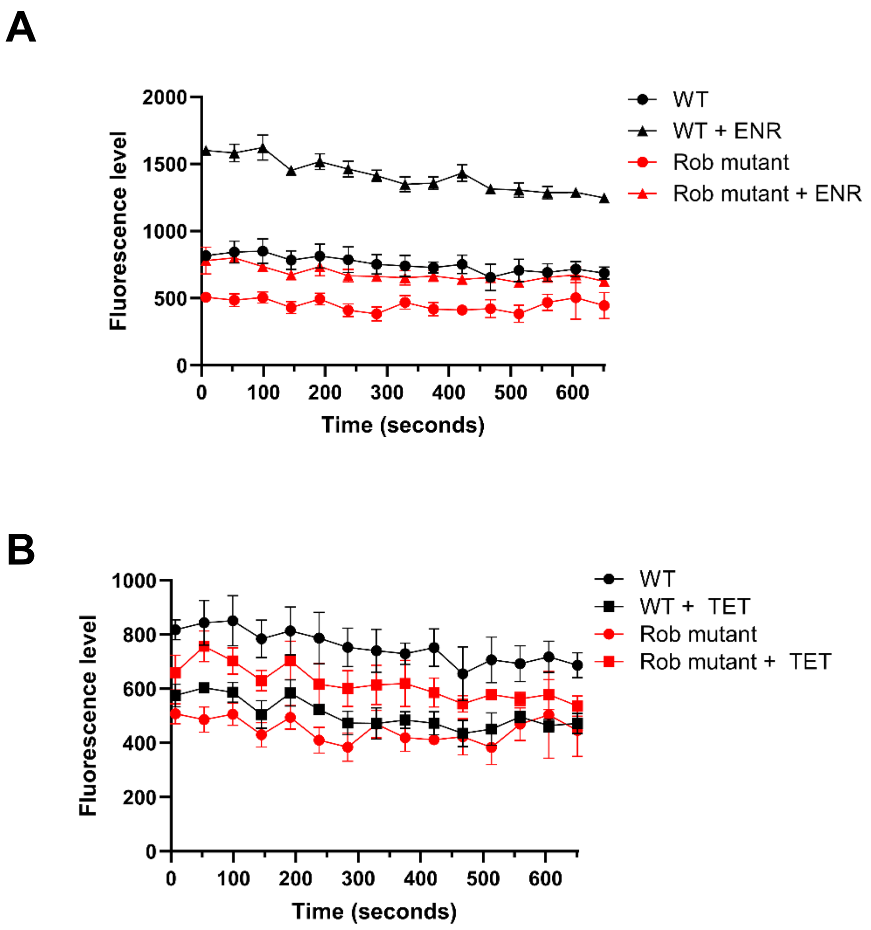


**Fig. S5. Nile Red efflux assay in wild-type and *rob* mutant strains**

Efflux activity quantified by residual Nile Red fluorescence under in the presence of **(A)** 1/16×MIC ENR; **(B)** 1/8×MIC TET. Time-resolved Nile Red efflux kinetics in wild-type and *rob* mutant strains. Cells were harvested from 23-h stationary-phase cultures, loaded with Nile Red, and efflux was initiated by the addition of 25 mM glucose at t = 0. Fluorescence was monitored continuously over time using excitation and emission wavelengths of 550 nm and 650 nm, respectively. Endpoint fluorescence values quantified at the final time point of the assay (cycle 15, approximately 11 min after glucose addition) were used for the statistical analyses shown. Data represent mean ± SD from three independent biological replicates with three technique replicates.

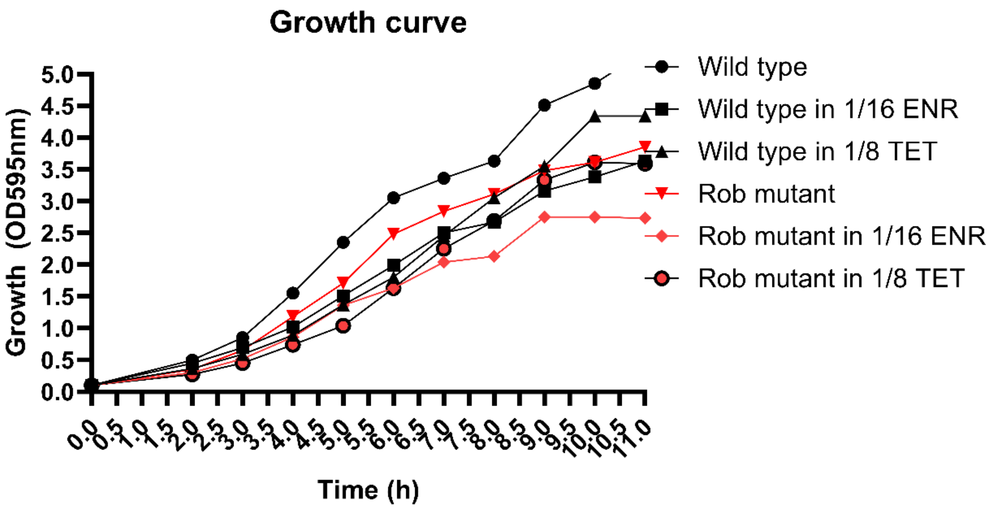


**Fig. S6. Measurement of growth for RNA sequencing in wild-type and *rob* mutant strains cultured in 20-mL flasks**

Growth curves of wild-type and *rob* mutant strains measured in 20 mL shake flasks over 11 h. Cultures were grown under drug-free conditions, in the presence of 1/16× MIC enrofloxacin (ENR), or 1/8× MIC tetracycline (TET). Growth was monitored by measuring optical density at 595 nm ($OD_{595}$) at the indicated time points. Growth rates were calculated to determine the early exponential phase at 3.5 h, starting from an $OD_{595}$ of 0.05. Data represent mean values from independent cultures from three independent biological replicates with three technique replicates.

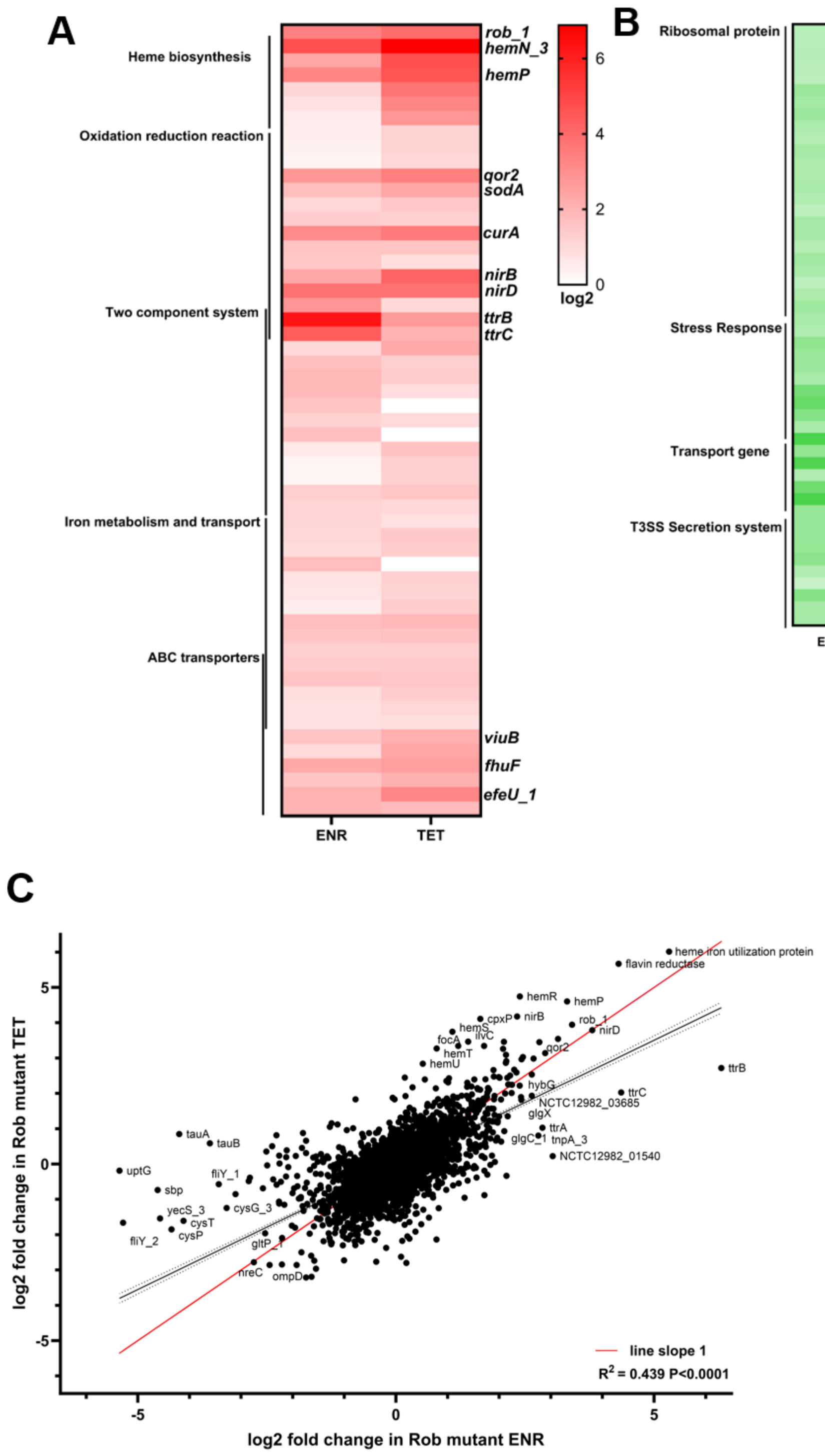


**Fig. S7. Transcriptional response of the *rob* mutant to sub-MIC antibiotic exposure.**

**(A)** Heatmap showing genes upregulated in the *rob* mutant relative to the wild type during growth in the presence of 1/16× MIC enrofloxacin (ENR) or 1/8× MIC tetracycline (TET). Genes are grouped by functional categories, including heme biosynthesis, redox reactions, two-component systems, iron metabolism and transport, and ABC transporters. Color scale indicates log2 fold change. **(B)** Heatmap of genes downregulated in the *rob* mutant under ENR and TET treatment, highlighting functional groups such as ribosomal proteins, stress response genes, transport genes, and the type III secretion system (T3SS). Color scale indicates log2 fold change. **(C)** Correlation analysis of genome-wide transcriptional changes in the *rob* mutant under ENR and TET exposure. Each dot represents an individual gene. The red line indicates a slope of 1. The coefficient of determination ($R^2 = 0.439$) and statistical significance ($P < 0.0001$) indicate a strong positive correlation between responses to the two antibiotics.

**A**

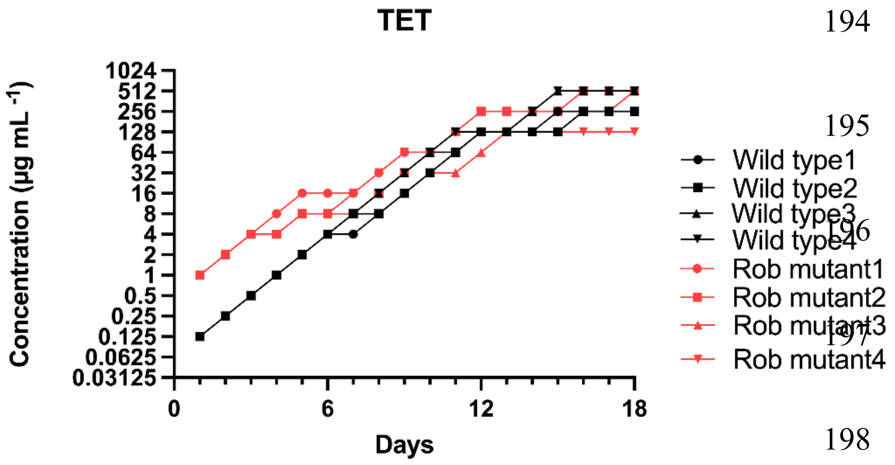


**B**

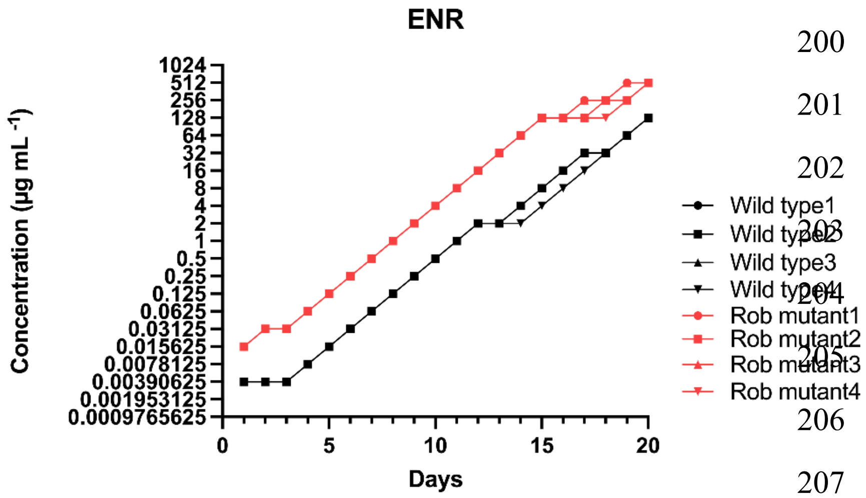


**Fig. S8. Resistance evolution of wild-type and *rob* mutant strains under tetracycline and enrofloxacin selection.**

**(A)** Step-wise increases in tetracycline (TET) concentration tolerated by four independent wild-type lineages (black) and four independent *rob* mutant lineages (red) during serial passaging. Cultures were propagated daily, and antibiotic concentrations were increased when populations showed stable growth. **(B)** Step-wise increases in enrofloxacin (ENR) concentration tolerated by four independent wild-type lineages (black) and four independent *rob* mutant lineages (red) during serial passaging. Antibiotic concentrations (μg $mL^{-1}$) are plotted on a log2 scale as a function of time (days). Data are shown from four independent biological replicates.

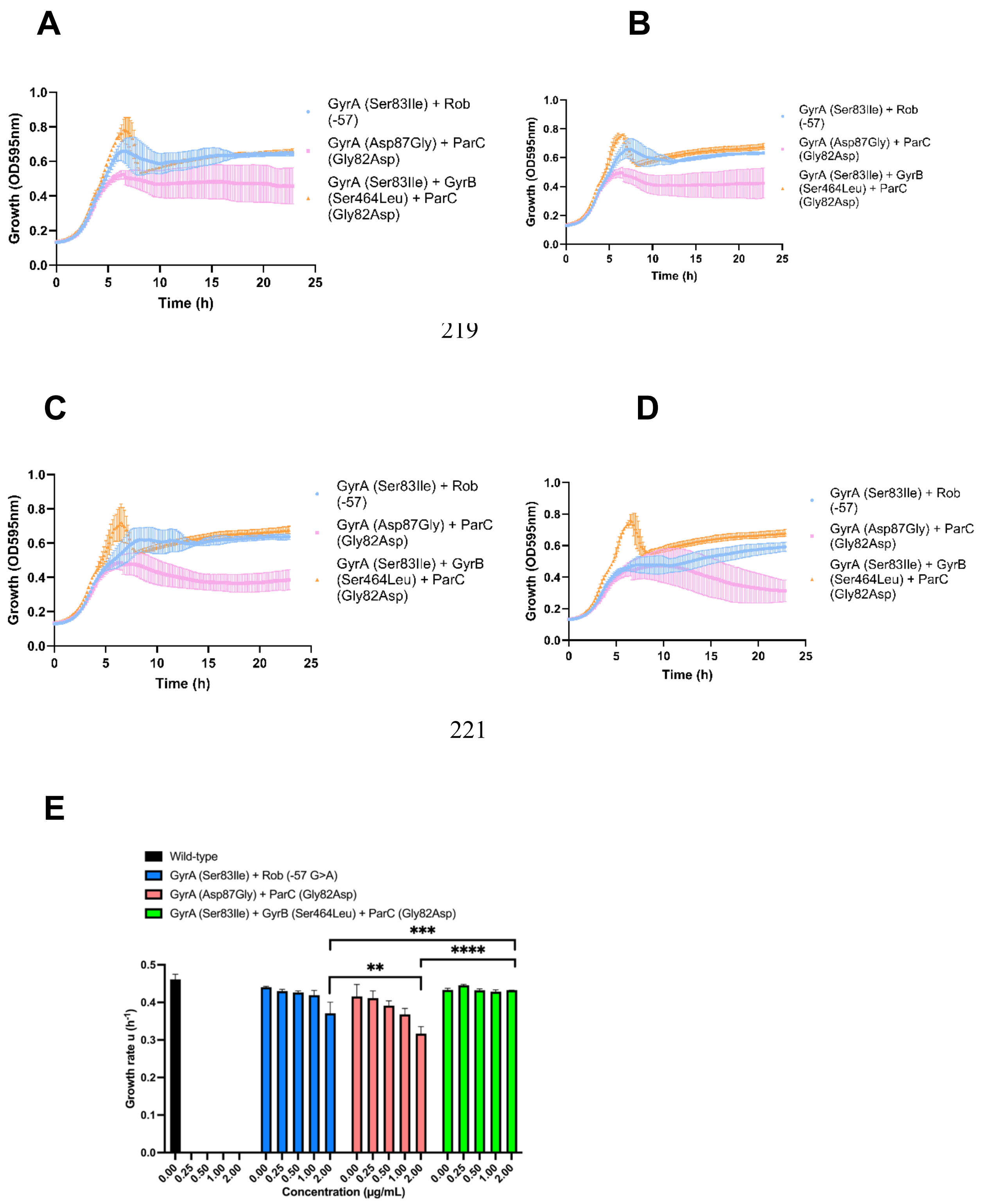


**Fig. S9. The growth of three mutational lineages under weak enrofloxacin exposure**.

**(A–D)** Growth curves of three mutational lineages measured under increasing enrofloxacin (ENR) concentrations (0.25, 0.5, 1, and 2 μg/mL). Curves represent mean growth ± SD over time. **(E)** Quantification of growth rates derived from panels

A–D across ENR concentrations. Bars represent mean growth rate (μ, $h^{-1}$) ± SD. Strains include wild-type (black), *gyrA* (Ser83Ile) + *rob* (-57 G>A) (blue), *gyrA* (Asp87Gly) + *parC* (Gly82Asp) (red), and *gyrA* (Ser83Ile) + *gyrB* (Ser464Leu) + *parC* (Gly82Asp) (green). Growth rates were comparable among the three lineages in the absence of ENR and at low ENR concentrations (0.25 and 0.5 μg/mL), indicating similar fitness and potential clonal interference under weak antibiotic selection. Statistical analyses were performed using two-way ANOVA with multiple-comparison correction; pairwise comparisons are indicated. **$P < 0.01$, ***$P < 0.001$, ****$P < 0.0001$.

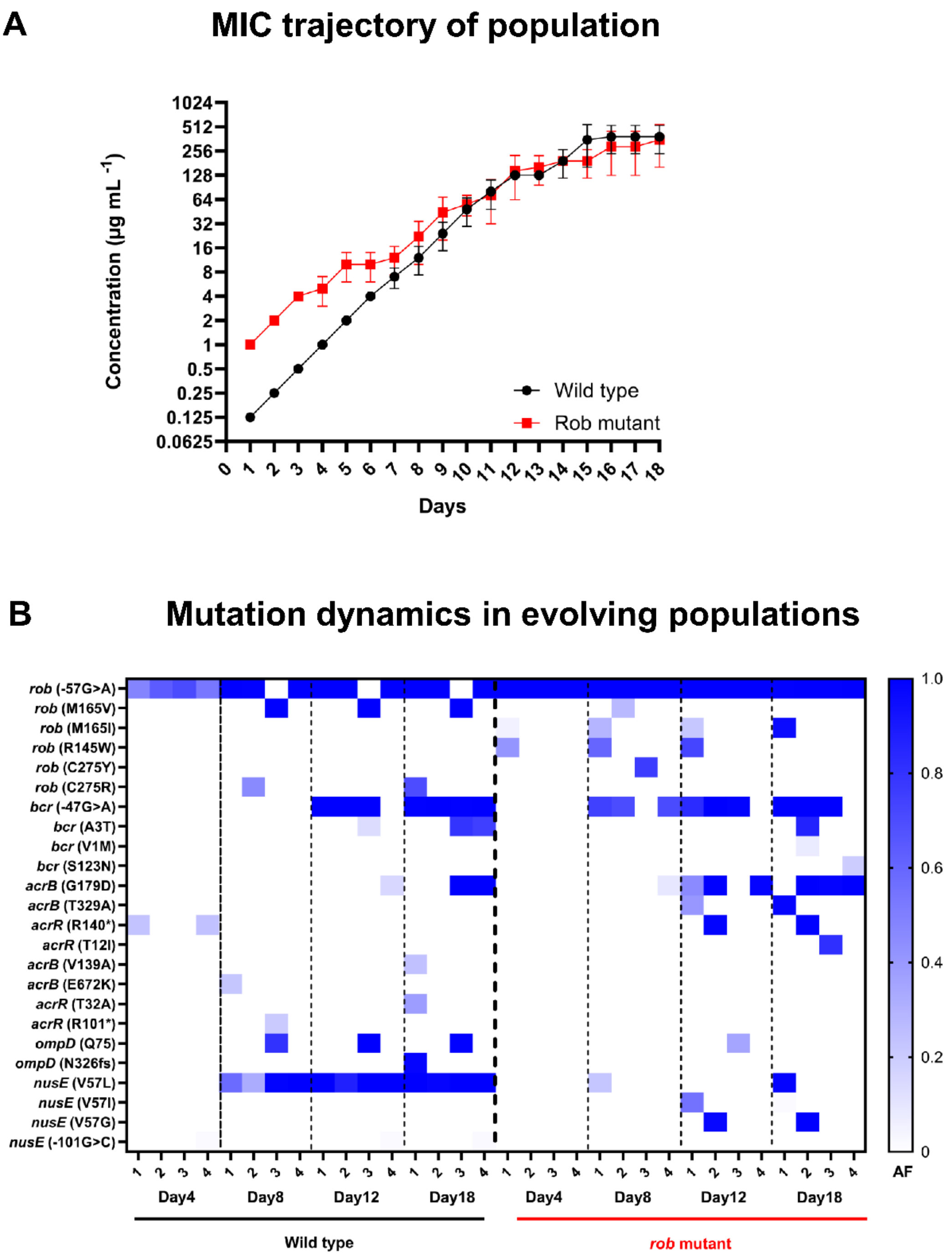


**Fig. S10. MIC trajectories and mutation dynamics during antibiotic-driven evolution of wild-type and *rob* mutant populations.**

**(A)** Population-level minimum inhibitory concentration (MIC) trajectories of wild-type (black) and *rob* mutant (red) populations during serial passaging under antibiotic selection. Points represent mean MIC values across replicate populations, and error bars indicate standard deviation. Antibiotic concentrations (μg $mL^{-1}$) are plotted on a log2 scale as a function of time (days). **(B)** Mutation dynamics in evolving wild-type and *rob* mutant populations. Heatmap shows the allele frequency (AF) of recurrent mutations detected by whole-population sequencing at the indicated time points. Each row represents a specific mutation, and each column corresponds to a sampling day. Blue color intensity indicates increasing allele frequency. Dashed vertical lines separate sampling intervals, and populations are grouped by genetic background (wild type vs *rob* mutant). Data are shown from four independent biological replicates.

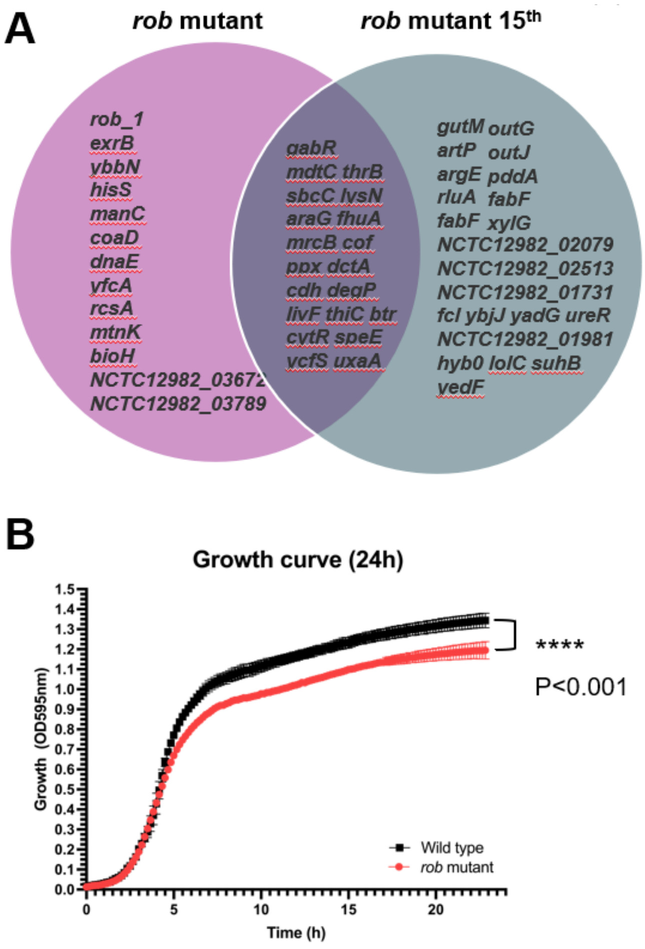


**Fig. S11. Genetic instability and fitness cost of the *rob* (-57 G>A) mutation.**

**(A)** Comparison of mutations introduced during strain construction with de novo mutations acquired in the *rob* mutant after 15 serial propagations (allele frequency > 0.5). **(B)** Growth curves over 24 h reveal a significant reduction in overall growth of the *rob* mutant compared with the wild type. Data represent mean ± SD from three technical replicates. Growth curves were compared using two-way repeated-measures ANOVA with multiple-comparison correction.

| ***rob* Mutant Construction** | **Sequence (5′→3′)** |
|---|---|
| PDS132-Vector-F Forward | TAGCGTACGCCATGGAGATCT |
| PDS132-Vector-R Reverse | TAGCTCGAGAGGGCCCTTAAGGTG |
| PDS132-Rob-vec.F: | ATCGAGCTCTCCCGGGAATTC |
| PDS132-Rob-vec.R: | ATCGCATGCGGTACCTCTAGAA |
| PDS132-Rob-Frag.F: | TTCTAGAGGTACCGCATGCGATCCAGCAGCATAATACTGCGGGT |
| PDS132-Rob-Frag.R: | GAATTCCCGGGAGAGCTCGATCCATGCAAGAGGAATGCTGCCCA |
| *rob*-F-SphI cut: | GGCATGCCCAGCAGCATAATACTGCGG |
| *rob*-R-SacI cut: | CCGAGCTCCCATGCAAGAGGAATGCTGC |
| *rob*-F-New | GTTATTATCCATCATCTGCGCGCTATTAAG |
| *rob*-R-New | AGCATTGCCTTGATGAGAAATCGCCAG |
| **Rob Flag Construction** | |
| R-FLAG-US | GTAATCGATGTCATGATCTTTATAATCACCGTCATGGTCTTTGTAG<br>TCGTTCACCGACATAAGGTAAGTGTAGG |
| F-FLAG-DS | GGTGATTATAAAGATCATGACATCGATTACAAGGATGACGATGAC<br>AAGTAACGCTCAACTGCGACGGCAAAA |
| F-ARM-US | AGAGGTACCGCATGCGATATACAGTTTTACTGGTGGGGATAATTAGGC |
| R-ARM-DS | TTCCCGGGAGAGCTCGATTGTGTAGATGCCTGTCCGGCG |
| AMPLIFY-PDS132-F | ATCGAGCTCTCCCGGGAA |
| AMPLIFY-PDS132-R | ATCGCATGCGGTACCTCT |
| **Colony Isolation** | |
| GyrA-F | CAA TCA ACC GCG CCT TGT TCA C |
| GyrA-R | AGG CAA CTG CCT CAA ACT TGG |
| GyrB-F | ACC GGC TCA AAC AGC TGA CG |
| GyrB-R | TGC GCG TAA AGC TCG TGA AAT G |
| ParC-F | AAC AGT AGT GCC ATT GAG CAG A |
| ParC-R | AGC CTG ATA CTG GCC AAC CTG |
| Rob-F | GGT ATT ATC CAT CAT CTG CGC GCT ATT AAG |
| Rob-R | AGC ATT GCC TTG ATG AGA AAT CGC CAG |
| **Gene Expression by qPCR** | |
| 16S rRNA-F | GCACGTAATGGTGGGAACTC |
| 16S rRNA-R: | CTCCAATCCGGACTACGACA |
| Rob-q-F | ATATACGTGGACGGGTACTGACG |
| Rob-q-R | TGCACAGCCCACTGTTCCATC |
| **Amplicon and Sanger Sequencing** | |
| Rob Forward | GTT ATT ATC CAT CAT CTG CGC GCT ATT AAG |
| Rob Reverse | AGC ATT GCC TTG ATG AGA AAT CGC CAG |

**Table S1** Primers used in sequencing and construction

| Gene | Mutation | M * | M 1 | M 2 | M 3 | M 4 | M 5 | M 6 | W 1 | W 2 | W 3 | W 4 | W 5 | W 6 | W 7 | W 8 |
|---|---|---|---|---|---|---|---|---|---|---|---|---|---|---|---|---|
| *ybbN* ← | D265G (GAT→GGT) | ✓ | | ✓ | ✓ | ✓ | ✓ | ✓ | | | | | | | | |
| ***ureR*** ← | coding (829/870 nt) | | | | | | | | ✓ | ✓ | ✓ | ✓ | ✓ | ✓ | ✓ | ✓ |
| ***rob_1*** ← / → ***nqrB*** | intergenic (-57/-433 | ✓ | | ✓ | ✓ | ✓ | ✓ | ✓ | | | | | | | | |
| *lolC* ← | S321S (AGC→AGT) | | | | | | | | ✓ | ✓ | ✓ | ✓ | ✓ | ✓ | ✓ | ✓ |
| *fabF_1* ← | C395R (TGT→CGT) | | | | | | | | ✓ | ✓ | ✓ | ✓ | ✓ | ✓ | ✓ | ✓ |
| *artP* → | Q239* (CAA→TAA) | | | | | | | | ✓ | ✓ | ✓ | ✓ | ✓ | ✓ | ✓ | ✓ |
| *aroC* ← | A39T (GCT→ACT) | | | | | | | | ✓ | | ✓ | ✓ | ✓ | ✓ | ✓ | ✓ |
| *sixA* ← / ← *fadJ* | intergenic (-327/+55)) | ✓ | | ✓ | ✓ | ✓ | ✓ | ✓ | | | | | | | | |
| *gutM* → | R84H (CGC→CAC) | | | | | | | | ✓ | ✓ | ✓ | ✓ | ✓ | ✓ | ✓ | ✓ |
| *suhB_2* → | S58S (TCT→TCC) | | | | | | | | ✓ | ✓ | ✓ | ✓ | ✓ | ✓ | ✓ | ✓ |
| *fabF_3* ← | P228P (CCT→CCC) | | | | | | | | ✓ | ✓ | ✓ | ✓ | ✓ | ✓ | ✓ | ✓ |
| *hyb0* ← | G340G (GGA→GGG) | | | | | | | | ✓ | ✓ | ✓ | ✓ | ✓ | ✓ | ✓ | ✓ |
| *bioH* → | Y5C (TAT→TGT) | ✓ | | ✓ | ✓ | ✓ | ✓ | ✓ | | | | | | | | |
| *xylG* ← | A501A (GCA→GCG) | | | | | | | | ✓ | ✓ | ✓ | ✓ | ✓ | ✓ | ✓ | ✓ |
| *coaD* ← | A136A (GCA→GCG) | ✓ | | ✓ | ✓ | ✓ | ✓ | ✓ | | | | | | | | |
| *yedF* → | S9N (AGT→AAT) | | | | | | | | ✓ | ✓ | ✓ | ✓ | ✓ | ✓ | ✓ | ✓ |

| | | | | | | | | | | | | | | | | |
|---|---|---|---|---|---|---|---|---|---|---|---|---|---|---|---|---|
| ***NCTC12982_03671 → / → NCTC12982_03672*** | **intergenic (+554 /-138)** | ✓ | | ✓ | ✓ | ✓ | ✓ | ✓ | | | | | | | | |
| ***NCTC12982_03789 →*** | **S41G (<u>A</u>GT→<u>G</u>GT)** | ✓ | ✓ | ✓ | ✓ | ✓ | ✓ | ✓ | | | | | | | | |
| ***yadG →*** | **F272S (T<u>T</u>C→T<u>C</u>C)** | | | | | | | | ✓ | ✓ | ✓ | ✓ | ✓ | ✓ | ✓ | ✓ |
| ***mtnK →*** | **R365W (<u>C</u>GG →<u>T</u>GG)** | ✓ | | ✓ | ✓ | ✓ | ✓ | ✓ | | | | | | | | |
| **tnp_1** | **(Δ712 bp)** | ✓ | | | | | | | | ✓ | | ✓ | ✓ | | ✓ | ✓ |

**Table S2** Genetic background of evolved and constructed *Y. enterocolitica* strains
**M***: Constructed mutant; **M**: mutants carrying the -57 G>A mutation in *rob*, isolated on day 5 or day 8; **W**: wild-type strains lacking the -57 G>A mutation in *rob*, isolated on day 5 or day 8.

**Rob-150 bp promoter sequence (5’ to 3’)**

ttcattgttctttaaatatataattacactgcatactacattaatctatcattatgccaacgcttttctcaactctcatttttcacgttaat
aaatcgttaacttacaattaattacaaataaaccacccacaatcatcgctattataat

**Clustal Omega analysis**

**>E.coli_K-12_MarA**

MSRRNTDAITIHSILDWIEDNLESPLSLEKVSERSGYSKWHLQRMFKKETGHS
LGQYIRSRKMTEIAQKLKESNEPILYLAERYGFESQQTLTRTFKNYFDVPPHKY
RMTNMQGESRFLHPLNHYNS

**>Y.enterocolitica_Rob_1**

MINEDILFIDELIEWIEINLEKRPTLDDVAKISGYSKWHLQRKFKSIVGLQLASY
IRGRVLTRAAVALRISRRPIIEISDELGFDSQQTFTRTFKKRFGVTPNSFRQMEQ
WAVQGMLPRFNFYENYTPEIKRVSLPAQELVGFTRQLNFDEHNHCSGQHSSC
MAMKDEILLDFFKEVNFSCQRVYSLFSADKDQQGQKSMYYSTAIDKEKRNEI
QGHREIDSISIPKGEFLAISHQGNAKECIKFSIYLFNEVLPKLRDEFGGGIEMEVI
EVDSCHAESKLRDITAAYTYLMSVN

**>S.enterica_LT2_RamA**

MTISAQVIDTIVEWIDDNLNQPLRIDDIARHAGYSKWHLQRLFMQYKGESLG
RYVRERKLKLAARDLLDTDQKVYDICLKYGFDSQQTFTRIFTRTFNLPPGAYR
KEKHGRTH

**>Kpn_NCTC13443_RamA**

MTISAQVIDTIVEWIDDNLHQPLRIDDIARHAGYSKWHLQRLFLQYKGESLGR
YIRERKLLLAARDLRDTDQRVYDICLKYGFDSQQTFTRVFTRTFNQPPGAYRK
ENHSRAH

**>E.coli_K12_SoxS**

MSHQKIIQDLIAWIDEHIDQPLNIDVVAKKSGYSKWYLQRMFRTVTHQTLGD
YIRQRRLLLAAVELRTTERPIFDIAMDLGYVSQQTFSRVFRRQFDRTPSDYRHR
L

**>Y.enterocolitica_Rob_2**

MDQASIIRDLLSWLESHLDQPLALDNVAAKAGYSKWHLQRMFKDVTGNAIG
AYIRARRLSKAAVALRLTSRPILDIALQYRFDSQQTFTRAFKKQFAQTPALYRR
AEDWHSAGICPPIRLGDYTLPQPEFITLPEQHLVGLTQSYSCTLEQIMTHHSELR
VHFWQQYLGDADHLPPVLYGLHHSRPNQEKDDEQEIFYTTAIEPQHVPGNVQ
EGQPVILQGGEYVQFSYDGPPDGLQDFILTLYGTCLPQLALTRRCGYDIERFFP
QGRPKEGPPASLKCEYLIPIRR

**>Y.pestis_Rob**

MDQASIIRDLLSWLESHLDQPLALDNVAAKAGYSKWHLQRMFKDVTGNAIG
AYIRARRLSKAAVALRLTSRPILDIALQYRFDSQQTFTRAFKKQFAQTPALYRR
AEDWHSSGICPPIRLGTYTLPQPEFITLPEQHLVGITQSYSCTLEQISTHRAELRL
HFWQQYLGDADQLPPVLYGLHHSRPNPEKDDEQEIFYTTAIEPQHIPCNVPEG
QPVILQGGEYVQFSYDGPLDGLQNFILTLYGTILPQLALIRRRGYDIERFYPQG
RPKDGPPATLKCDYFIPIRR

**>Y.pseudotuberculosis_Rob**

MDQASIIRDLLSWLESHLDQPLALDNVAAKAGYSKWHLQRMFKDVTGNAIG
AYIRARRLSKAAVALRLTSRPILDIALQYRFDSQQTFTRAFKKQFAQTPALYRR
AEDWHSSGICPPIRLGTYTLPQPEFITLPEQHLVGITQSYSCTLEQISTHRAELRL
HFWQQYLGGADQLPPILYGLHHSRPNPEKDDEQEIFYTTAIEPQHVPCNVPEG
QPVILQGGEYVQFTYDGPLDGLQNFILTYLDTILPQLALIRRRDYDIERFYPQG
RPKDGPPATLKCDYFIPIRR

**>Kpn_295R_Rob**

MDQAGIIRDLLSWLEGHLDQPLSLDNVAAKAGYSKWHLQRMFKDVTGHAIG
AYIRARRLSKSAVALRLTARPILDIALQYRFDSQQTFTRAFKKQFSLTPALYRRS
PDWSSFGMRPPLRLGEFTLPKHEFITRPPTQLLGVTQSYTCKLEEISDFRNQMR
VQFWRDFLGNSPSIPPVLYGLHEPRPSLEKDDEQEVFYTTALTPEMANGHLQH
AHPVTLEGGEYVMFTYEGLGTGLQEFILTVYGTCMPMLNLTRRKGLDIERFYP
EDESRDQATPIQLRCEYLIPIRR

**>Ecloacae_E1252_Rob**

MDQAGIIRDLLTWLEGHLDQPLSLDNVAAKAGYSKWHLQRMFKDVTGHAIG
AYIRARRLSKSAVALRLTARPILDIALQYRFDSQQTFTRAFKKQFSLTPALYRRS
PDWSSFGMRPPLRLGEFAMPKYDFITLPETHLIGTTQSYSCSLEQISEFRHQMR
VQFWRDFLSHAPAIPPILYGLNETHPSQEKDDEQEVFYTTALTPEMANGYIQGS
KPVVLEGGEYVMFAYEGLGTGVQEFILTVYGTCMPMLNLNRRKGQDIERYYP
AQDAKPEEGPINLRMEFLIPIRR

**> S.enterica_LT2_Rob**

MDQAGIIRDLLIWLEGHLDQPLSLDNVAAKAGYSKWHLQRMFKDVTGHAIG
AYIRARRLSKSAVALRLTARPILDIALQYRFDSQQTFTRAFKKQFSQTPALYRRS
SEWSAFGIRPPLRLGEFTVPEHQFVTLEDTPLLGVTQSYSCSLEQISDFRHEMR
VQFWHDFLGHSPTIPPVLYGLNETRPSMEKDDEQEVFYTTALPQEQADGYVQ
SAHPVLLQGGEYVMFTYEGLGTGVQDFILTVYGTCMPMLNLTRRKGQDIERY
YPSEDTKTGDRPINLRCEFLIPIRR

**>Citrobacter freundii_ICC168_Rob**

MDQAGIIRDLLVWLEGHLDQPLSLDNVAAKAGYSKWHLQRMFKDVTGHAIG
AYIRARRLSKSAVALRLTARPILDIALQYRFDSQQTFTRAFKKQFAQTPALYRRS
PEWSAFGIRPPLRLGEFAIPEHKFVTLEDTQLVGITQSYSCSLEQISDFRHEMRV
QFWHDFLGHAPAIPPVLYGLNETRPSLEKDDEQEVFYTTALTPEQANGYVQTA

QPVLLQGGEYVMFTYEGLGTGVQEFILTVYGTCMPMLNLTRRKGQDIERYYP
AEDAKAGDRPINLRCEFLIPIRR

**>E.coli_K12_Rob**

MDQAGIIRDLLIWLEGHLDQPLSLDNVAAKAGYSKWHLQRMFKDVTGHAIG
AYIRARRLSKSAVALRLTARPILDIALQYRFDSQQTFTRAFKKQFAQTPALYRRS
PEWSAFGIRPPLRLGEFTMPEHKFVTLEDTPLIGVTQSYSCSLEQISDFRHEMR
YQFWHDFLGNAPTIPPVLYGLNETRPSQDKDDEQEVFYTTALAQDQADGYVL
TGHPVMLQGGEYVMFTYEGLGTGVQEFILTVYGTCMPMLNLTRRKGQDIER
YYPAEDAKAGDRPINLRCELLIPIRR

**>Sflexneri_301Ser2a_Rob**

MDQAGIIRDLLIWLEGHLDQPLSLDNVAAKAGYSKWHLQRMFKDVTGHAIG
AYIRARRLSKSAVALRLTARPILDIALQYRFDSQQTFTRAFKKQFAQTPALYRRS
PEWSAFGIRPPLRLGEFTMPEHKFVTLEDTPLIGVTQSYSCSLEQISDFRHEMR
YQFWHDFLGNAPTIPPVLYGLNETRPSQDKDDEQEVFYTTALAQDQADGYVL
TGHPVMLQGGEYVMFTYEGLGTGVQEFILTVYGTCMPMLNLTRRKGQDIER
YYPAEDAKAGDRPINLR
